# Archaeogenetic analysis revealed East Eurasian paternal origin to the Aba royal family of Hungary

**DOI:** 10.1101/2024.03.20.585718

**Authors:** Gergely I B Varga, Zoltán Maróti, Oszkár Schütz, Kitti Maár, Emil Nyerki, Balázs Tihanyi, Orsolya Váradi, Alexandra Ginguta, Bence Kovács, Petra Kiss, Monika Dosztig, Zsolt Gallina, Tibor Török, János B. Szabó, Miklós Makoldi, Endre Neparáczki

**Affiliations:** Department of Archaeogenetics, Institute of Hungarian Research, Budapest, Hungary; Department of Pediatrics and Pediatric Health Center, University of Szeged, H-6725 Szeged, Hungary; Department of Genetics, Institute of Biology, University of Szeged, H-6726 Szeged, Hungary; Department of Biological Anthropology, University of Szeged, H-6726 Szeged, Hungary; Department of Archaeology, Institute of Hungarian Research, Budapest, Hungary; Department of History, Institute of Hungarian Research, Budapest, Hungary

## Abstract

The Aba family played a pivotal role in the early history of Medieval Hungary dominating extensive territories and giving rise to influential figures. We conducted an archaeogenetic examination of remains uncovered at the necropolis in Abasár, the political centre of the Aba clan, to identify Aba family members and shed light on their genetic origins. Utilizing Whole Genome Sequencing (WGS) data from 19 individuals, complemented by radiocarbon measurements, we identified 6 members of the Aba family who shared close kinship relations. Our analysis revealed that 4 males from this family carried identical N1a1a1a1a4∼ haplogroups. Significantly, our phylogenetic investigation traced this royal paternal lineage back to Mongolia, strongly suggesting its migration to the Carpathian Basin with the conquering Hungarians. Genome analysis, incorporating ADMIXTURE, Principal Component Analysis (PCA), and qpAdm, revealed East Eurasian patterns in the studied genomes, consistent with our phylogenetic results. Shared Identity by Descent (IBD) analysis confirmed the family kinship relations and shed light on further external kinship connections. It revealed that members of the Aba family were related to members of prominent Hungarian medieval noble families the Árpáds, Báthorys and Corvinus as well as to the first-generation immigrant elite of the Hungarian conquest.

## Introduction

Over the past few decades, the collaboration between archaeology and archaeogenetics has opened up new avenues for identifying the remains of renowned historical figures and exploring their familial history. Such combined methods enabled the identification of Richard III’s skeleton (*1*), the remains of Romanov family members (*2*), or Birger Magnusson, the founder of Stockholm (*3*). This approach was also the key to identify the famous Hungarian king, Matthias Corvinus’ descendants (*4*), or the members of the Báthory family, one of the most prominent aristocratic families of Medieval Central Europe (*5*). Archaeogenetic methods have also facilitated the identification of members of the Árpád dynasty, Hungary’s first royal family, and to analyse their genomic heritage (*6*–*9*). Hence, archaeogenetics has become a valuable method for addressing both prehistoric and historic inquiries.

The Abas were one of the most prominent Hungarian noble families, holding extensive territories in Heves County (Northern Hungary) during the Middle Ages. The ethnic origin of the Aba family is a subject of controversy in historical records (*10*). According to the first surviving Hungarian medieval historical text Anonymus’ Gesta, Ed and Edemen, the progenitors of the clan, were identified as Cuman chiefs who had aligned themselves with the tribal alliance of the conquering Hungarians in present-day Russia prior to the Hungarian Conquest. As a result, they were granted extensive territories in Northern Hungary by Prince

Árpád (*10*–*13*) Since we know that the Cumans only moved to this area in the mid-11th century, it seems that Anonymus’ text anachronistically referred to the population, which living here before the arrival of the people of Almos and Árpád as Cuman (*14*). According to the 13^th^-century historian Simon of Kéza and later the 14^th^-century Chronicon Pictum, the forefathers of the clan, Ed and Edemen were the sons of Csaba, who was the legendary son of Attila, the grand King of Huns, according to medieval Hungarian historical tradition (*12, 13, 15, 16*). The latest historical work, Chorincon Pictum was the first to claim that the Abas and the Árpáds would have had a direct common ancestor, Attila’s son Csaba (*12, 16*). Modern historians propose an Eastern origin for the family, as both the Cumans and Huns can trace their origins back to the East. The most plausible ancestral groups linked to the Aba lineage are the Kabar clans, who separated from the Khazar Khaganate and joined the Hungarians shortly before the conquest (*12, 17*).

The honoured progenitor of the clan, Samuel Aba (Sámuel in Hungarian) (c. 990-1044), initially served as the comes of the court (*comes palatii*) to Hungary’s first Christian king, St. Stephen I (István I) (ruled 1000-1038). Sámuel was the “sororius” of the first King. One possible explanation is that this meant that due to his elevated status, entered into marriage with one of the king’s sisters (*12*). The other possibility is that Samuel was the king’s sister’s son, so nephew (*18*). Thus, through this marriage and their descendants, the Abas established a kin relationship with the Árpád dynasty. Sámuel later ascended to the throne as the third monarch of the Kingdom of Hungary (ruled 1041-1044), becoming the country’s first elected king. It is assumed that Sámuel’s social and political advancement may have been influenced by his family’s presumed prominent noble origin.

Following the death of István I, the power passed to his nephew, Peter Orseolo (Péter I) (ruled 1038-1041 and 1044-1046). Péter ruled with a firm hand, appointing foreign nobles to key positions, which elicited discontent among the Hungarian populace. Consequently, a revolt erupted in 1041, resulting in Péter’s deposition and the subsequent enthronement of Sámuel Aba, who wielded significant influence among the rebels. However, his reign was short-lived. After three years of battling external and internal adversaries, he met his demise in the Battle of Ménfő in 1044, where he confronted claimant Péter, supported by the German monarch Henrik III (*12, 19, 20*).

King Sámuel’s body was interred temporarily until the completion of the church he founded in Abasár – the political hub of the clan in the late 10^th^ and early 11^th^ centuries. Subsequently, a few years later, his remains were exhumed and laid to rest in the Abasár church (*10, 12, 20, 21*). Despite Sámuel’s downfall, the family’s presence endured on the medieval Hungarian political landscape. While historical records from the 11th and 12th centuries lack information about the family, from the 13th century onwards, their names appear regularly in written sources. By this time, the clan had splintered into dozens of families, some of which possessed extensive territories within the kingdom. They wielded significant political influence as dignitaries and even oligarchs during the 13th to 15th centuries (*22*–*24*).

It is presumed that Abasár retained its position as an important political centre for the clan and served as the final resting place for numerous descendants. According to the Hungarian chronicles the monastery in Abasár was funded by Sámuel Aba in the first half of the 11^th^ century, however it was first documented in a diploma issued by king Béla IV in 1261 (*12, 25*). In later centuries, the monastery and its domains became the subject of numerous ownership disputes among different branches of the clan, including the Nánai Kompolti, Csobánka and Ugrai families, and other Hungarian nobilities (*25*). By the end of the 14^th^ century the significance of the monastery had increased as the abbot of Abasár was mentioned twelve times in the papal bulls of Popes Boniface IX and Gregory XII (*25*). At the end of the 16^th^ century, the Nyáry family owned the possessions of the abbey, then during the Ottoman occupation those fell into the hands of the white clergy (*25*). In the 17^th^ century, the domains were embroiled in several ownership disputes, and the abbey subsequently vanished from the sources, decaying over time (*26*).

Between 2020 and 2022, the Department of Archaeology at the Institute of Hungarian Research conducted excavations at the Abasár Bolt-tető site in Northern Hungary, unveiling the remains of the church, primarily established by Sámuel Aba (*12, 25*) (Figure 1A-B). The excavation revealed various phases of the church’s existence, providing evidence of constructions, renovations, and instances of destruction. Although the tomb of the king could not be identified, graves belonging to notable members of the Aba family were discovered within the church building. In the sanctuary, an unearthed tomb with a stone cover (Figure 1. C) featured a carved depiction of the Aba family’s coat of arms (Figure 1. D) (for more details of the archaeological examination see Supplementary text). The inscription on the border of the cover stone revealed that János and Mihály, two individuals from the Aba clan, were laid to rest there during the early years of the 15^th^ century. Different branches of the Abas did rule extensive areas in Heves County in the 13^th^-14^th^ centuries (*23*), and according to the medieval diplomas the monastery belonged to at least three different branches of the clan, as mentioned earlier (*25*). Thus, it is not clear which branch they exactly belonged to. Additionally, a double grave was found at the geometrical centre of the church building, with its stone covers also adorned with the clan’s coat of arms (Figure 1. F). Two more graves were uncovered within the sanctuary (Figure 1. C), bringing the total count to at least five prominent burials that clearly belonged to significant family members. As in these graves multiple skeletons were discovered in multiple layers, in some cases it was hard to identify the prominent individuals with archaeological methods alone. For more details of the archaeological findings, see Supplementary texts.

**Figure 1.**
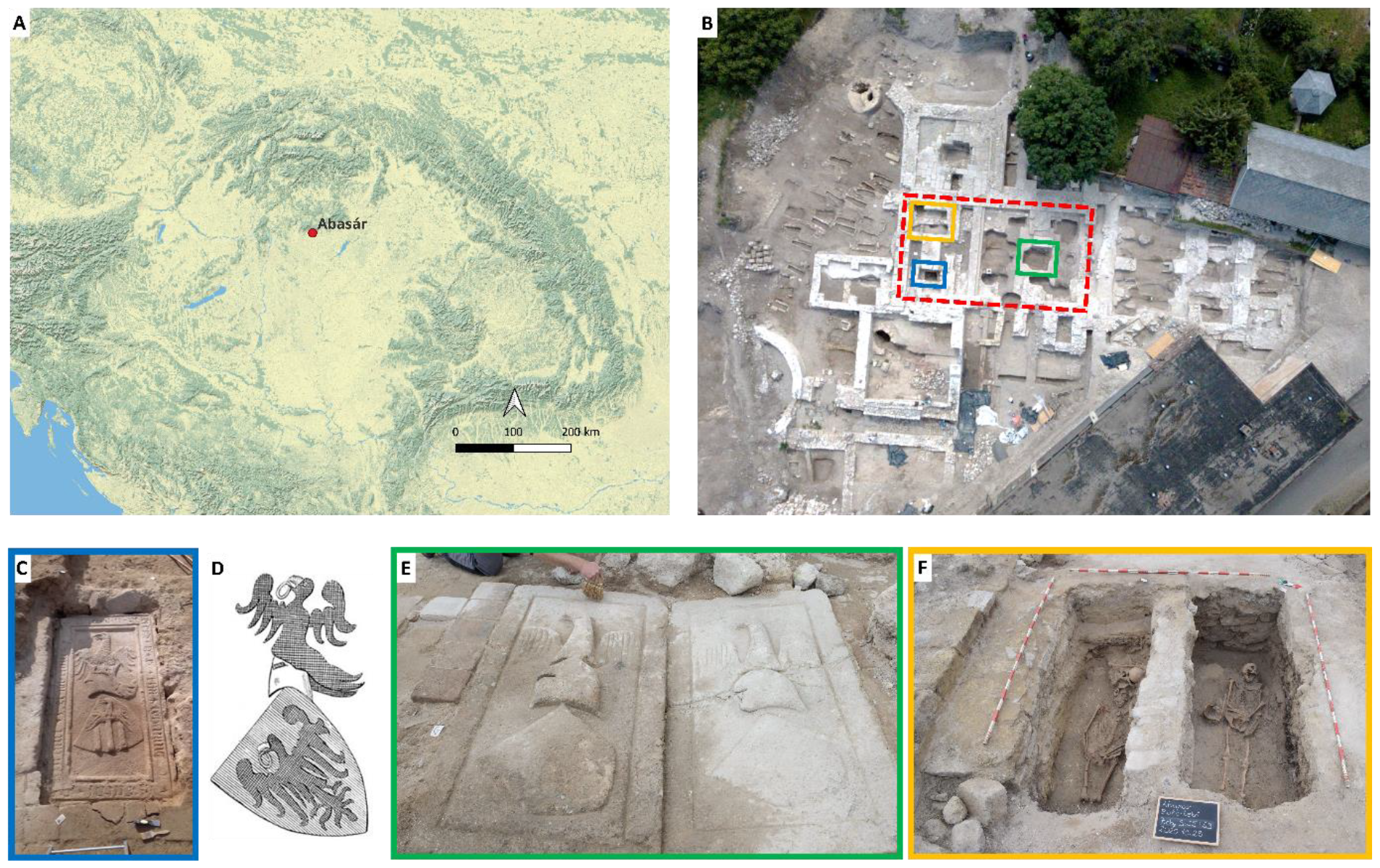
Abasár Bolt-tető site. A) The location of Abasár in Northern Hungary. B) The Abasár Bolt-tető site and the ruins of the monastery of Abasár viewed from above. The silhouette of the huge church building with the excavated graves is well discernible. The sanctuary is marked with a red dashed quadrate, while the exact positions of the prominent graves are indicated with yellow, blue and green quadrates respectively. C) The prominent burial in the sanctuary, at the altar district with the carved and scripted cover stone (HUAS57 and HUAS581) (Blue). The inscription indicates that outstanding members of the Aba family were buried in the grave. D) The stylistic depiction of the Abas’ coat of arms (27) which can be observed on the cover stones.E) The double grave in the geometrical centre of the church Phase I (HUAS261 and HUAS262) (green). F) The graves of HUAS55B and HUAS59B individuals in the altar district (yellow). The position of the burials refers to the prominence of the deceased inside.

Although the royal tomb and the king’s remains could not be identified, the archaeogenetic analysis of the interred individuals inside the church provided an unparalleled opportunity to explore the genetic origins of one of medieval Hungary’s most influential dynasties. As the paternal lineage of the Árpáds had previously been identified (*6*–*9*), we also had the opportunity to determine whether they shared the same paternal lineage as the Abas.

In this study, we present the results of our archaeogenetic investigation, employing a combination of archaeological and genetic methodologies to identify the family members, establish their kin relations, and analyse the phylogenetic connections of the Abas’ paternal lineage. Furthermore, we employ state-of-the-art genome analysis techniques to delineate their ancestral heritage.

## Results

### The Medieval genomic dataset from Abasár

Owing to medieval customs, kings and prominent figures were interred within the confines of churches. Therefore, we collected samples from all the skulls excavated within the edifice. Multiple burials were discovered in the prominent graves within the sanctuary and at the geometric centre of the church, but through careful analysis, we were able to identify the primary graves of potential members of the Aba family. These prominent remains are referred as HUAS55B, HUAS57, HUAS581, HUAS59B, HUAS261, and HUAS262 throughout the study. For a comprehensive account of the excavation and archaeological discoveries, please refer to the Supplementary Materials.

DNA was successfully extracted from the total of 38 indoor remains from Abasár. Double-stranded sequencing libraries were constructed, incorporating partial UDG treatment (refer to Table S1). The libraries from 20 Abasár samples met the quality criteria and were sequenced to achieve a 1.6x average genome coverage (0.5-3.11x). The observed *post mortem* damage (PMD) ratio aligned with the general patterns of ancient DNA. Rigorous testing for contamination in mitochondrial DNA and heterozygosity of polymorphic sites on the X chromosome in males revealed a minimal level of contamination in our dataset. Applying the genetic sex estimation method of Skoglund et al. 2013 (*28*), two remains were identified as genetically female, while the remaining samples, were conclusively identified as males (Table S1). More detailed information on DNA extraction, library preparation, sequencing, PMD, contamination and sex determination, is also provided in Table S1.

Radiocarbon analysis of ten skeletons dated the Abasár site to the period ranging from the 12th to 15th century CE (Table 1, Supplementary table 1 and Supplementary Text3). This analysis corroborated historical and preliminary archaeological data (Supplementary text1 and 2), confirming that the burials can be attributed to the era of the medieval Kingdom of Hungary. Specifically, three notable individuals from stone-covered graves were dated from the late 13th to the end of the 14th century, closely matching the approximate date engraved on the stone cover indicating the burials took place in the very early years of the 15th century.

**Table 1.** Radiocarbon data of the studied remains. All Abasár samples were dated to the era of the medieval Kingdom of Hungary, corresponding to historical data.

|  | SAMPLE TYPE | CAL. CE |
| --- | --- | --- |
| HUAS261 | tooth | 1277-1301 (88,4%); 1371-1378 (7%) |
| HUAS262 | tooth | 1278-1301 (87,8%); 1371-1378 (7,6%) |
| HUAS309 | tooth | 1305-1365 (76,6%); 1383-1398 (18,9%) |
| HUAS341 | tooth | 1455-1505 (78,8%); 1596-1618 (16,6%) |
| HUAS390 | tooth | 1169-1222 (95,4%) |
| HUAS450 | tooth | 1301-1329 (43,8%); 1341-1369 (29,3%); 1379-1396 (22,3%) |
| HUAS55B | rib | 1276-1300 (92,6%); 1372-1377 (2,8%) |
| HUAS57 | tooth | 1279-1303 (79%); 1368-1379 (16,4%) |
| HUAS59B | rib | 1300-1325 (43,7%); 1352-1395 (51,7%) |

### Kinship analysis

As the Abasár Bolt-tető site was believed to be the family cemetery of the Aba clan, we anticipated discovering kin relationships as well as identical uniparental haplotypes among the individuals under investigation.

The kinship analysis using correctKin revealed several family connections further supported by the uniparental data. Two small families were identified from the prominent graves mentioned earlier: one comprising four individuals (HUAS55B, HUAS59B, HUAS57, and HUAS581) and another consisting of two individuals (HUAS261 and HUAS262) (see Table S2).

The family of four, including one female and three males, was excavated from the sanctuary of the church. Two males, HUAS57 and HUAS581, turned out to be 5^th^ degree relatives. Notably, these were uncovered from the grave with the carved stone cover (Figure 1C) of which inscription indicates that two significant male members of the clan were buried there. Two other members of the same family, HUAS55B male and HUAS59B female, were excavated from two uncovered tombs in the sanctuary (Figure 1F) and were found to be first-degree relatives of each-other with identical Mt haplotype. The HUAS59B female was a 3^rd^ degree relative of HUAS57 and 4^th^ degree relative of HUAS581 from the inscribed tomb, while the HUAS55B male was 4^th^ degree relative of HUAS57 and its relation to HUAS581 could not be determined, as it was beyond 5th degree. HUAS55B and HUAS59B obviously were in a mother-son relationship, also confirmed by IBD analysis, as they shared 3477 centiMorgan (cM), the entire length of the genome with each-other (Table S7). For the reconstructed plausible family trees of this family refer to Extended Figure 1.

The other two related male individuals (HUAS261 and HUAS262), found in the prominent stone-covered double grave in the geometric centre of the church (Figure 1E), and were determined to be 3rd degree relatives. It is noteworthy that all four males unearthed from decorated tombs with cover stones shared the same N1a1a1a1a4∼ Y-chromosome haplogroup. suggesting that two branches of the same extended family or clan were identified in this cemetery. Subsequent IBD analysis confirmed that the two families were indeed two branches of the same extended family.

### Phylogenetic connections of the Aba paternal lineage

Out of 5 males in the extended Aba family 4 carried identical N1a1a1a1a4∼ Hgs. Our comprehensive analysis revealed a diverse phylogenetic network of Hg N1a1a1a1a4∼, and its sub-branches (Figure 2). Utilizing the Yleaf software with the markers of ISOGG 2020, the Aba lineage could be classified within the N1a1a1a1a4a2∼ (N-A9408) sub-branch, except for the sample HUAS57, which lacked coverage over the A9408 marker of the Hg. Notably, two elite conquering Hungarians, along with an unpublished elite Xiongnu (AG6F) from the Ar Gunt site, Mongolia, also shared this sub-Hg. This suggests a Mongolian origin for the lineage that arrived in the Carpathian basin with the conquerors.

**Figure 2.**
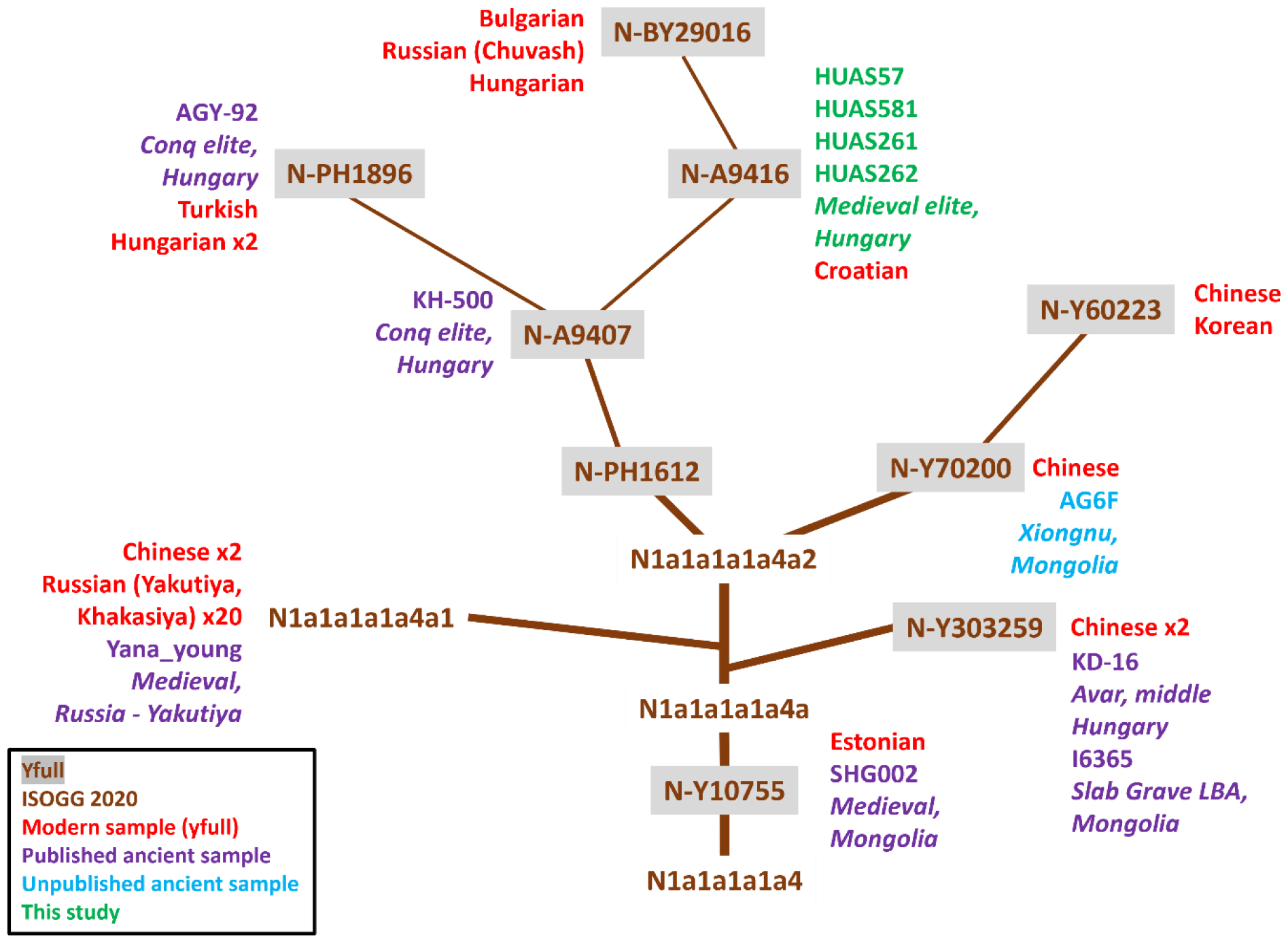
The phylogenetic analysis of Hg N1a1a1a1a4. The tree was generated according to the data of ISOGG2020 and yfull.com. Hg assignment of ancient sequences was conducted based on SNPs collected from each ancient genomes in the Y chromosome positions indicated in Table S3. The studied lineage belongs to the N-A9416 subbranch near the branch of elite members of the conquering Hungarians.

Enhancing our analysis with markers and modern data from the yfull database enabled us to position the ancient samples within the deeper branches of the phylogenetic tree. The N-A9408 Hg bifurcates into Eastern and Western branches. The Eastern branch, N-Y70200, is present in modern-day Chinese and Korean individuals, with the unpublished Xiongnu sample allocated to this sub-branch. The Western branch, N-PH1612, is predominantly found in Europeans, and after the Hg N-A9407 it further divides into two sub-branches: N-PH1896 and N-A9416. The conqueror samples belong to N-PH1896 and N-A9407, along with a contemporary Turkish individual and two Hungarians. The Aba lineage, along with a modern Croatian male, was assigned to N-A9416. The sub-Hg of N-A9416 is also present in a Hungarian, a Chuvash, and a Bulgarian individual.

The initial identification of Hg N1a1a1a1a4∼ traces back to a Mongolian individual from the Late Bronze Age Slab Grave culture (*29*). Presently, it exhibits the highest prevalence among Yakut males and is also notably common among Evenks and Evens (*30*). These findings strongly indicates an origin in Inner Asia. Our phylogenetic tree further supports this narrative, revealing that its sub-branch, N-PH1612, made its way to Europe and Hungary through medieval migrations including the conquering Hungarians.

### Uniparental haplogroups and phylogenetic connections of the cemetery

The mitochondrial genomes from Abasár showed high heterogeneity, with 18 different haplotypes determined (Table S1c). HUAS55B and HUAS59B belonged to the same R0b haplotype, consistent with the result of the kinship analysis. Most of the mitochondrial lineages (14/18) have been detected in preceding populations of the Carpathian basin (*31*–*33*) and several were present in the late medieval Hungarian elite cemetery of the Báthory family (*5*). Based on the comparison with our published ancient whole mitogenome database (*31*), eight lineages have a feasible Near Eastern (H13a1d, R0b and T2) and/or Steppe origin (H28, T1a1, I1b, HV14a and J1c5a), while the remainder are most probably of European origin.

As anticipated in the case of a family cemetery, the Y chromosomal Hgs displayed much less diversity. Among the 17 males there were representatives from 13 different sub-branches of the Y chromosomal phylogenetic tree of ISOGG (https://isogg.org/tree/index.html). In addition to individuals belonging to the royal family with Hg N1a1a1a1a4∼, different I1∼, I2∼, R1a1a1b1∼, R1a1a1b2∼ and R1b∼ lineages were represented by single individuals, while one pair belonged to Hg R1a1a1b1a2b3a3a2g2∼ (Table S1c). Similarly, to the mitochondrial lineages, most of the Y-chromosomal lineages (12/17) had European origin. A considerable number (7/17) were detected in preceding periods of the Carpathian basin, and several were prevalent among contemporary Hungarian elite (*5, 32*). Besides the royal lineage, only one Hg with obvious Asian origin was detected: R1a1a1b2a2a3c2∼. It belongs to the Asian subbranch of R1a, specifically R1a-Z2125, which emerged in the Middle-Late Bronze Age among people of the Sintashta and Andronovo cultures. It was widespread among the Scythians and their descended populations (*29, 34*–*36*). The sub-Hg found in Abasár has been detected in Hun samples (*37*), and its supergroups were prevalent among Huns and Avars in the Carpathian basin (*32*). It has also been found among the Pazyryk and Kangju peoples (*38, 39*). Today, R1a1a1b2a2a3c2∼ is primarily observed in males from Eastern and Central Eurasia, with a higher prevalence in Russian Tatars.

### Genomic heritage of the Abasár remains

The analysis of uniparental markers revealed an Inner-East Asian paternal origin for the Aba clan, raising the question of whether these ancestral ties were also discernible within their genomes. To address this question, we conducted genome analyses, leveraging the power of ADMIXTURE, principal component analysis (PCA), and qpAdm.

The ADMIXTURE analysis (K=7) unveiled a strikingly similar genome component pattern between the Abasár group and Germanic populations from early medieval Germany and Slovakia. Additionally, Hungarian samples from commoner (village) cemeteries of the conquering period exhibited a remarkably comparable genetic profile too (Figure 3 and Table S4). The Abasár genomes were composed of 0-3,5% Nganasan-, 6-11% Western Hunter-Gatherer-, 38-45% European neolithic farmer-, 0-3,5% She-, 0-9% Iranian neolithic-, 3,5-13% Early Bronze Age Western Eurasian- and 29-39% Ancient North Eurasian (ANE)-related genome components. The East Eurasian (Nganasan, She) components indicate the presence of minor eastern ancestry.

**Figure 3.**
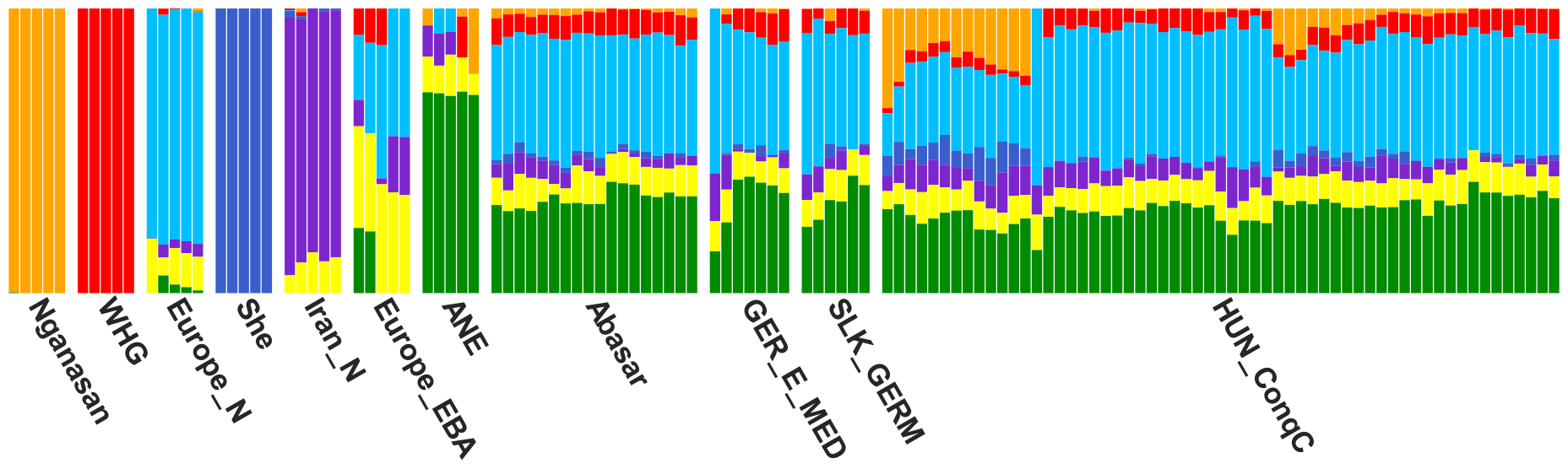
Unsupervised ADMIXTURE analysis (K=7) results of the Abasár individuals. Besides the early medieval population from Bavaria, Germany and a Germanic group from modern-day Slovakia, the commoner people from the Hungarian conquering era of the Carpathian basin had the most similar genome composition to the Abasár samples.

On the European PCA (Figure 4 and Extended Figure 2), the majority of the samples clustered together, separate from the cloud of Central European populations, shifted to the direction of the conquering Hungarians, with some outliers falling within the variance of present-day Hungarians. The Abasár cluster overlaps with the genomes of the Árpád dynasty, as well as with the two Báthorys, a prominent dynasty of late medieval Hungary and Poland (*5*). Notably, the Abasár cluster also overlaps with the cluster of other samples from the cemetery of the Báthory family at Pericei (Extended Figure 3). As Abasár Bolt-tető site and the Pericei graveyard are thought to be aristocratic cemeteries, this observation suggests a characteristic and uniform genome composition for the medieval Hungarian noble stratum. On the Eurasian PCA, the Abasár group also exhibited a noticeable eastward genetic shift compared to modern Europeans (Extended Figure 4). Moreover, most samples overlapped with the genetic cline of the conquering Hungarians from the 9-11th centuries (Figure 4 and Extended Figure 4). Both ADMIXTURE and PCA findings underscored the eastern connections of the Abasár genomes, potentially linked to the subsequent influx of steppe immigrants into the Carpathian basin.

**Figure 4.**
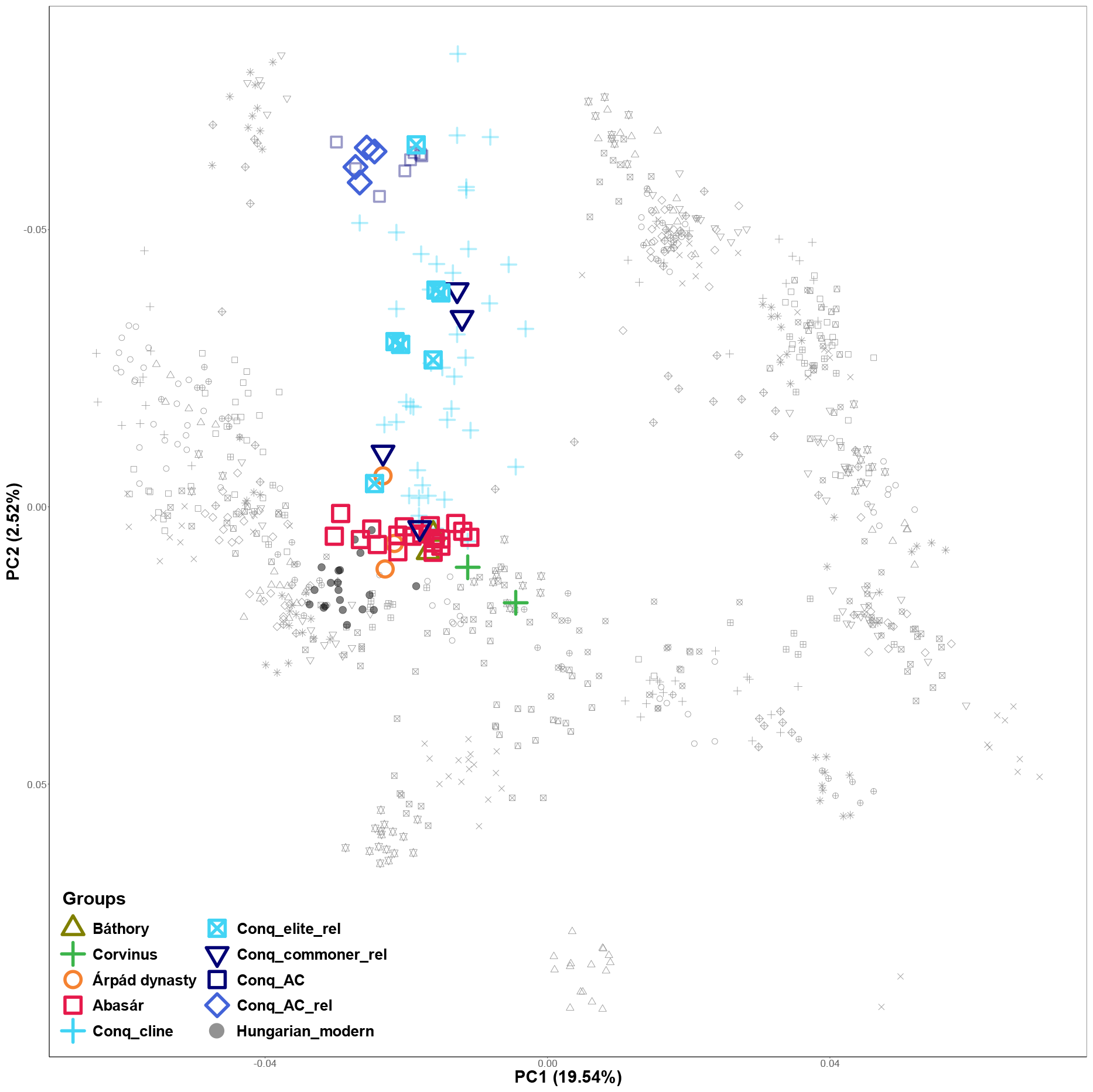
European PCA of the Abasár genomes (red squares) projected onto the axes determined by 784 modern genomes (light gray shapes). The studied samples form a distinct cluster from modern-day Central European peoples overlapping with the cline of the conquering Hungarians (skyblue crosses), and some outliers fall into the variance of present-day Hungarians (dark grey spots). The Abasár samples mapped close to the remains of the Báthorys (greenish brown triangles), the Corvins (green crosses) and the Árpád dynasty members (gold circles). The immigrant core group of the conquerors is also included (Conq_AC, dark blue squares). Conq_elite_rel (blue square with X-shape), Conq_commoner_rel (dark blue triangle) and Conq_AC_rel (blue diamonds) represent the conquerors sharing significant length of IBDs with the Abas based on our IBD sharing analysis (see below).

To identify the plausible source of the Asian genome elements in the Abasár samples, we conducted qpAdm analysis. In the left population list, we included contemporaneous and preceding Medieval populations from the Carpathian basin and Central Europe, a medieval group from Northern Caucasus (Anapa), and eastern immigrant groups of the Carpathian basin from the Middle Ages. The ‘base model strategy’ run resulted in dozens of plausible two-source models for almost all samples (Table S6, summarized in Figure 5).

**Figure 5.**
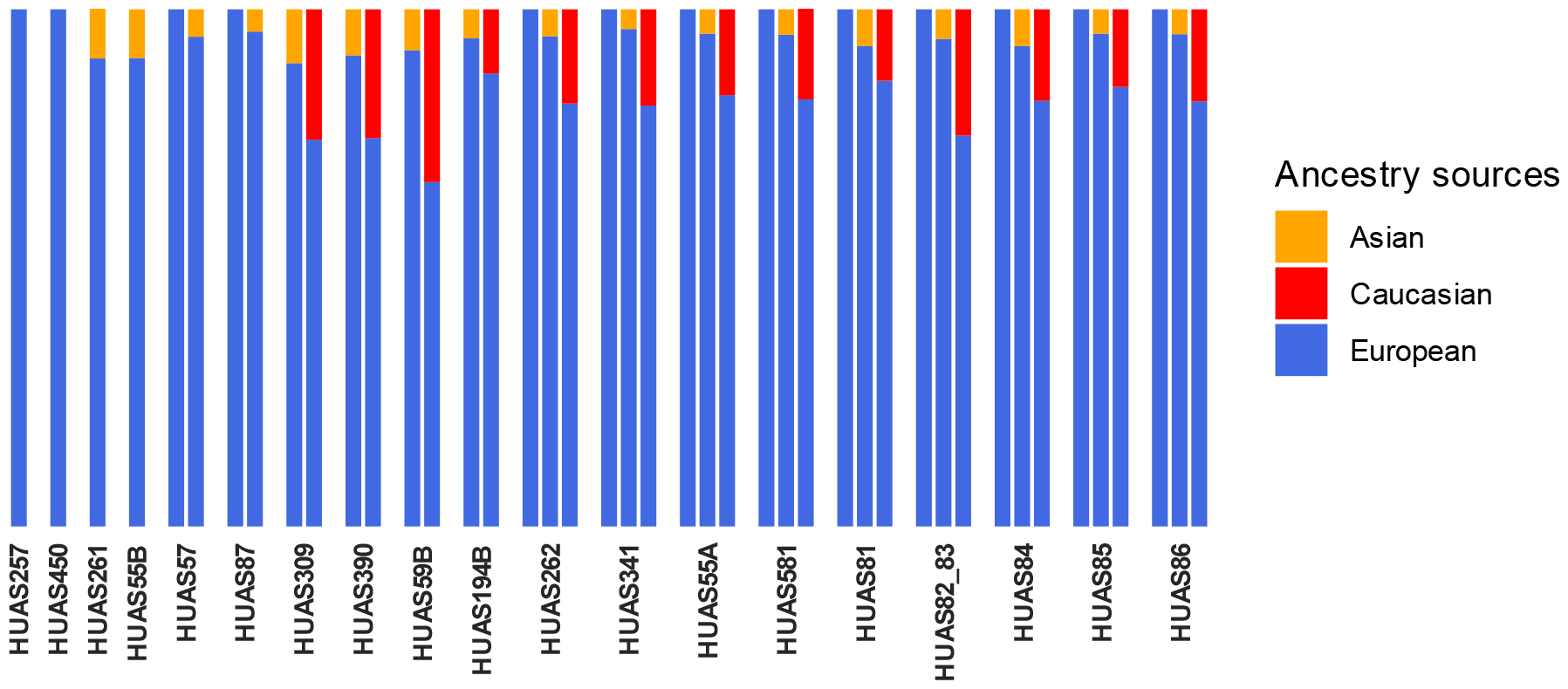
qpAdm models for the medieval genomes from Abasár Bolt-tető site with significant p-values. In most cases multiple models were valid with different European, Asian and Caucasian source combinations indicating a low amount of Asian ancestry in the Abasár genomes.

Two samples (HUAS257 and HUAS450) could be modelled from single European sources, forming clades with Germanic or Carpathian basin sources, and two could be obviously modelled as an admixture of major European and minor Asian sources. In all other samples, qpAdm unequivocally indicated the admixture of European and different Eastern sources subserving the results of ADMIXTURE and PCA. Equivalent models of these genomes with comparable p-values supported the single-source explanation or identified significant minor components of Asian (Hun/Avar/conquering Hungarian) and/or Caucasian origin. The average ratio of the Asian ancestry in these genomes ranged between 4,3-10,4%.

### Shared IBD analysis

In order to identify genetic relations of the Abasár individuals at a finer scale, we utilized genome imputation and conducted IBD analysis using ancIBD (Table S7). We identified shared IBD segments longer that 8cM between the Abasár genomes and those of the Huns, Avars and conquering Hungarians published by (*32*). We also tested IBD sharing with the additional medieval samples from Hungary, as the Pericei genomes (*5*) the Corvinus (*4*) and the Árpáds (*8*). In the IBD graph of Figure 6, we illustrated all detected IBD connections between the samples from Abasár and other samples with a minimum cumulative IBD length of 10 cM.

**Figure 6.**
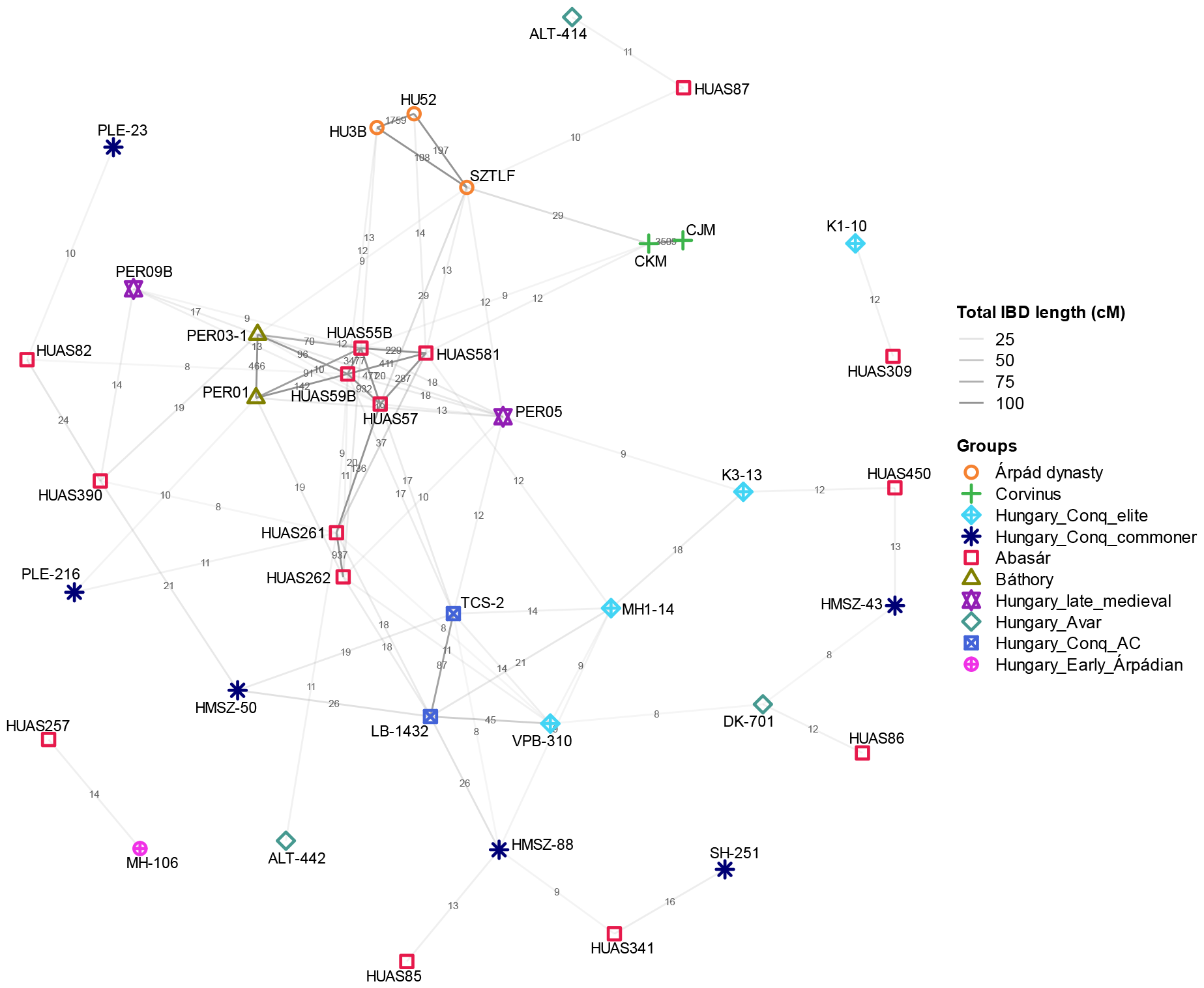
Shared IBD network. The network indicates the IBD connections among the Abasár remains and other medieval individuals from the Carpathian basin, including the elite individuals of the conquering Hungarians, and the members of different aristocratic families as the Corvinus, the Báthorys and the Árpáds. The length of the edges linking the individuals is inversely proportional with the total length of shared IBDs between the two genomes. Total IBD length is indicated at the middle of each edge. The network was generated in R, with the packages igraph and ggplot2.

First of all, the IBD graph distinctly illustrates the familial patterns, with cumulative lengths of shared IBD among family pairs closely aligning with their genealogical relatedness as measured by correctKin. Notably, this analysis revealed a significant amount of shared IBD among members of the two distinct families, indicating an approximately 6-7th degree of relationship between them. This emphasizes the identification of two branches within a single extended family or clan. Furthermore, we identified two additional Abasár remains (HUAS390 and HUAS82) as distant relatives to the two families, suggesting that members of distantly related branches of the clan were also interred within the church. This observation aligns with historical data, which documents at least three branches of the clan governing the monastery during the 13^th^-15^th^ centuries.

Importantly, based on shared IBDs, these families exhibit distant connections to medieval and early modern Hungarian noble lineages, including the Corvins (*4*) and the Báthorys (*5*). The shared IBDs between some Abasár samples and members of the Árpád dynasty align with historical records, which document at least one marriage between the two families. Additionally, various samples excavated in Abasár share IBDs with different noble individuals but not with the aforementioned Aba family, suggesting their affiliation with other noble families of the Kingdom of Hungary.

The IBD connections to the immigrant core of the Hungarian conquering elite (*32*) imply that the genetic relationship of the family to the conquerors predates the settlement of the Hungarians in the Carpathian basin. This conclusion aligns with the results of our phylogenetic investigation and definitively establishes that the minor eastern genomic component originates from the conquering Hungarians.

## Discussion

Medieval Hungarian kings, including Sámuel Aba, were traditionally interred in churches, following the custom of Christian monarchs. While nearly half of the medieval Hungarian rulers found their final resting place in Székesfehérvár (*40, 41*), within the coronating basilica founded by István I, the others were buried in temples that they had either privately funded or renovated for funeral purposes. Unfortunately, the ravages of time lead to the destruction of these temples, and the tombs of many kings were lost. Béla III’s intact burial discovered in 1848 in Székesfehérvár was a rare exception (*42, 43*). According to Hungarian chronicles, Sámuel Aba was buried in the monastery of Abasár, a foundation attributed to him. However, the church associated with his burial site vanished over the centuries, and the Abasár Bolt-tető site was identified as the most likely location (*25*). The archaeological findings of the new excavation provided further evidence confirming that this was the monastery of the king.

In the stone graves inside the church, human remains were uncovered in primary anatomical order. Some tombs contained multiple skeletons arranged in both primary and secondary positions, indicating the reuse of graves across successive generations. Instances of an ossuary-like arrangement of skeletons indicated episodes where numerous remains were exhumed and then collectively reburied in common graves, possibly during reconstruction, rebuilding operations, grave robberies, or mass grave burials due to epidemics or wars. The part of the destruction is likely associated with the Mongol invasion of 1241-42, as the site is located along the marching route of Tatar troops, who were known to devastate Christian churches and plunder graves. Therefore, the archaeological investigation encountered difficulties in pinpointing the royal burial of Sámuel Aba.

Despite the inability to locate the king’s tomb, the excavations unearthed burials of prominent family members inside the church, underscoring the enduring significance of the church in the clan’s history. A double grave and a single tomb, all adorned with stone covers depicting the coat of arms of the Aba clan were discovered. The sanctuary tomb, with an inscription indicating the interment of János and Mihály, members of the genus, in the 15th century, aligned with historical records, that large territories in Heves county, including the monastery of Abasár, were possessed by different branches of the clan Aba, including the Kompolti, Ugrai and Csobánka branches (*22, 23, 25*). Four males from these prominent graves shared the same Y chromosome Hg, likely representing two branches of the Aba family. The results of shared IBD analysis confirmed that two branches of one large family were uncovered in these tombs, and they obviously were nobilities as they had extended kinship network with other Hungarian noble families. Thus, archaeological and archaeogenetic data indicate that Aba clan members were identified at Abasár Bolt-tető site, and these results fit with the historical records.

Our IBD analysis has revealed dynastic connections among medieval Hungarian noble families, both those known from written sources and those previously unknown. The relationship through marriage was indicated in historical records between the Abas and the Árpáds (*10, 12, 19*), but we lack data in the relation of the Abas and the Báthorys, and also between the Abas and the Corvins. Nevertheless, we detected genealogical connections among all these prominent Hungarian families, indicating the existence of frequent marital relations in the noble stratum of the medieval Kingdom of Hungary. The results also revealed direct connection between the Abas and the first generation immigrant core of the conquering Hungarians (*32*), suggesting that this connection predates the Conquest Period. It suggests that the forefathers of the Abas were the members of the wondering and conquering Hungarian tribes, aligning with the results of the phylogenetic connections as well.

Regarding the family’s ethnic roots, historical sources have proposed Hunnic and Cuman origins, but the prevailing consensus among historians tends to favour Khazar or Kabar ancestry (*12, 17*), as well as direct conqueror nobility (*10*). Our phylogenetic results reveal that the paternal lineage of the family belongs to Hg N1a1a1a1a4∼, with Inner-Asian origin. This aligns with potential connections to the Hunnic, Cuman, Kabar or Khazar groups. However, the more deeply characterized sub-Hg of the Aba paternal lineage, N-A9416, belongs to the N-PH1612 subgroup identified in Eastern Europe, with the majority of carriers located in present-day Hungary. Among them the oldest samples include two elite individuals from the conquering Hungarians. Notably, two additional conqueror samples with Hg N1a1a1a1a4∼ were identified (*44*). However, later they were excluded from our analysis because the Hg determination of these samples relied on a hybridization-based amplicon sequencing method, which lacked information about inner SNP markers. Taking all the available data into account, it is highly probable that the paternal lineage of the Abas is associated with an elite paternal line among the conquering Hungarians. The paternal ancestors of the Aba family could potentially include one of the high ranked persons of the conquering Hungarian tribes.

In Hungarian chronicles both the Abas and the Árpáds are identified as descendants of Attila, the grand prince of the Hunnic empire. This would imply that the paternal lineages of the two families are the same. Since the R1a paternal lineage of the Árpáds has been previously identified (*6*–*8*) and it does not match the paternal lineage identified in the Abas, we can now definitively exclude this possibility. Nonetheless, the unverified assertion of either royal lineage being descended from Attila persists. Detailed Whole Genome Sequencing (WGS) data revealed the Árpáds’ paternal ancestry originating from East Eurasia (*6*), with potential Hunnic connections (*45*). In this study, the Abas also demonstrated Hunnic/Xiongnu phylogenetic connections, rendering both families to be credible candidate for this esteemed genealogy.

The genomic-level analyses align with the uniparental data. Both ADMIXTURE and PCA analyses indicated minor Asian ancestry in the studied medieval individuals. The qpAdm analysis also identified a notable minor East Asian component in the majority of the Abasár genomes; however, due to the low fraction of this component, the program could not pinpoint its precise source. Potential sources include the Huns, Avars, or conquering Hungarians, however, the IBD results clearly indicate that the minor Asian component among the above originated from the conquering Hungarians. In summary, all genetic results consistently point towards the Abas being descendants of one of the tribal elites among the conquering Hungarians.

## Materials and Methods

### Radiocarbon dating

Radiocarbon analysis was performed on the sampled bone fragments to confirm the archaeological dating of the remains. The measurements were done by accelerator mass spectrometry (AMS) in the AMS laboratory of the Institute for Nuclear Research, Hungarian Academy of Sciences, Debrecen, Hungary (AMS Lab ID: DeA-37107; technical details concerning the sample preparation and measurement: Molnár et al. 2013 (*46*)). The conventional radiocarbon date was calibrated with the OxCal 4.4.4 software (https://c14.arch.ox.ac.uk/oxcal/OxCal.html, date of calibration: 9th of January 2024) with IntCal 20 settings (*47*).

### DNA extraction, library preparation and sequencing

All 38 in-door remains from Abasár with available petrous bone, or tooth were undergone genetic sampling (Table S1a). All steps of sampling, DNA extraction and library preparation were carried out as described in the Supplementary materials of Varga *et al*., 2023 (*8*), in the joint, dedicated ancient DNA laboratory of the Department of Archaeogenetics, Institute of Hungarian Research and the Department of Genetics, University of Szeged. The double stranded libraries were shallow shotgun sequenced on Illumina iSeq platform to monitor their human DNA content (Table S1b). The selected libraries were sequenced deeper on NovaSeq platform to an average genome coverage of 1x (Table S1c-e).

### Data processing and quality assessment of the ancient sequences

The adapters of paired-end reads were trimmed with the Cutadapt software (*48*), and sequences shorter than 25 nucleotides were removed. Read quality was assessed with FastQC (*49*).The raw reads were aligned to GRCh37 (hs37d5) reference genome using the Burrow-Wheels-Aligner (v 0.7.17) software, with the MEM command in paired mode, with default parameters and disabled reseeding. Only properly paired primary alignments with ≥ 90% identity to reference were considered in all downstream analyses to remove exogenous DNA. Samtools v1.1 was used for merging the sequences for different lanes, sorting, and indexing binary alignment map (BAM) files (*50*). PCR duplicates were marked using Picard Tools MarkDuplicates v 2.21.3 (*51*). To randomly exclude overlapping portions of paired-end reads and to mitigate potential random pseudo haploidization bias, we applied the mergeReads task with the options “updateQuality mergingMethod=keepRandomRead” from the ATLAS package (*52*). Single nucleotide polymorphisms (SNPs) were called using the ANGSD software package (version: 0.931-10-g09a0fc5) (*53*) with the “-doHaploCall 1 -doCounts 1” options and restricting the genotyping with the “-sites” option to the genomic positions of the 1240K panel.

Ancient DNA damage patterns were assessed using MapDamage 2.0 (*54*) and read quality scores were modified with the Rescale option to account for post-mortem damage. Mitochondrial genome contamination was estimated using Schmutzi algorithm (*55*). Contamination for the male samples was assessed by the ANGSD X chromosome contamination estimation method (*56*), with the “-r X:5000000-154900000 -doCounts 1 - iCounts 1 -minMapQ 30 -minQ 20 -setMinDepth 2” options (Table S1).

The raw nucleotide sequence data of the samples were deposited to the European Nucleotide Archive (http://www.ebi.ac.uk/ena) under accession number: PRJEB72247.

### Sex determination, haplogroup assignment and kinship estimation

Biological sex was assessed with the method described in (*28*). Fragment length of paired-end data and average genome coverages (all, X, Y, mitochondrial) was assessed by the ATLAS software package (*52*) using the BAMDiagnostics task. Detailed coverage distribution of autosomal, X, Y, mitochondrial chromosomes was calculated by the mosdepth software (*57*) (Table S1b).

Mitochondrial haplogroup (Mt Hg) determination was performed with the HaploGrep 2 (version 2.1.25) software (*58*), using the consensus endogen fasta files resulting from the Schmutzi Bayesian algorithm (Table S1d). The Y Hg assessment was performed with the Yleaf software tool (*59*), updated with the ISOGG2020 Y tree dataset (Table S1c).

Kinship analysis was performed with correctKin (*60*) (Table S2). As reference population we applied the same database as in Varga et al 2023 (*8*).

### Comparative analysis of uniparental haplogroups

To shed light on the most feasible origin of the uniparental lineages of our samples, we compared them to ancient and modern databases. Mt Hgs were set against the modern- and ancient whole mitogenome databases published in Maár et al., 2021 (*31*). The Y chromosomal Hgs were analysed in virtue of the archaic dataset of the Allen Ancient DNA Resource (AADR) (*61*) and the modern database of yfull (www.yfull.com). The determination of the most plausible phylogenetic origin was based on whether the samples shared the same or the closest Hg with the studied samples.

### Y chromosomal phylogenetic analysis of the Aba family

For detailed phylogenetic analysis the bam files of all the available ancient Y chromosome sequences from the literature belonging to Hg N1a1a1a1a4 or its sub-haplogroups (sub-Hgs) were collected. The ancient database was supplemented with the Y chromosome sequence of an unpublished Xiongnu individual (AG6F) from Ar Gunt site, Mongolia (list of the ancient samples is given in Table S3). The previously published Y Hgs were reassessed with the Yleaf software tool (*59*), updated with the ISOGG2020 Y tree dataset, and all Hgdefining markers were collected and verified manually using the Integrated Genome Viewer (IGV) (*62*). Then, based on the marker set and phylogenetic tree of yfull (yfull.com/tree), all ancient and studied samples were classified into the deeper branches of the tree through manual examination of the given markers (Table S3). Phylogenetic tree was drawn based on the verified markers of the ancient samples, and the dataset was supplemented with the modern individuals from the yfull database confirming their classification (Table S3).

### Unsupervised ADMIXTURE

Unsupervised ADMIXTURE was carried out as described in (*32*). We applied 4003 genomes including 1314 modern (HO dataset) and 2689 ancient genomes with 18 genomes from this study, excluding all 1^st^ and 2^nd^ degree relatives from each dataset (Table S4). Accordingly, HUAS55B was also excluded from the analysis due to its 1^st^ degree kin relationship to HUAS59B.

### PCA

We used the modified modern West Eurasian genome data of (*63*) (Table S5A), with a total of 803 individuals, and the Eurasian genome data published in (*32*) confined to the HO dataset, to draw a modern PCA background on which ancient samples could be projected. However, in order to obtain the best separation of our samples in the PC1-PC2 dimensions, South-East Asian and Near Eastern populations were left out, and generally just 10 individuals were selected from each of the remaining populations, leaving 1397 modern individuals from 179 modern populations in the analysis (Table S5B). PCA Eigen vectors were calculated from the 803 and 1397 pseudo-haploidized modern genomes with smartpca (EIGENSOFT version 7.2.1) (*64*). All ancient genomes were projected on the modern background with the ‘‘lsqproject: YES and inbreed: YES’’ options. Since the ancient samples were projected, we used a more relaxed genotyping threshold (>50k genotyped markers) to exclude samples only where the results could be questionable due to low coverage.

### qpAdm

We used qpAdm (*65*) from the ADMIXTOOLS software package (*66*) for modelling our genomes as admixtures of two source populations and estimating ancestry proportions. The qpAdm analysis was done with the HO dataset, as in many cases suitable Right or Left populations were only available in this dataset. As possible European source populations we selected Migration Period populations from the Carpathian basin and Central Europe from the AADR, based on preliminary outgroupf3 and qpAdm data of the Abasár samples. We also included Huns, Avars, conquering Hungarians and Anapa groups from (*32*) as possible sources of Asian and Caucasian ancestries. For the list of Left and Right populations see Table S6.

During ‘base model strategy’ run we set the details:YES parameter to evaluate Z-scores for the goodness of fit of the model (estimated with a Block Jackknife). As we wished to identify the possible source of the minor Asian-like genome proportion of some Abasár genomes, we ran the analysis just with source combinations of two. As qpWave is integrated in qpAdm, the nested p values in the log files indicate the optimal rank of the model. This means that if p value for the nested model is above 0.05, the Rank-1 model should be considered (*65*).

### IBD sharing analysis

For imputation, we used the GLIMPSE2 framework (version 2.0.0) (*67*) using the 1KG Phase 3 dataset common markers as reference. The reference data set was normalized and multi allelic sites were split using bcftools (version 1.16-63-gc021478 using htslib 1.16-24-ge88e343) with the “norm –m –any” subcommand and filtered for biallelic SNPs with the “view –m 2 –M 2 –v snps” subcommand. The autosomal chromosomes of the human reference genome were divided into 580 genomic chunks using the GLIMPSE2_chunk tool with the “-sequential” option. As described in the GLIMPSE2 manuscript, we created the binary reference data with the GLIMPSE2_split_reference tool using the 580 genomic regions and the 1KG biallelic SNP variants.

In all downstream imputation analysis, we used only samples with >0.5x mean genome coverage of shotgun WGS data as recommended in the GLIMPSE2 manuscript. Furthermore, we excluded all samples with estimated MT contamination higher than 0.03 (based on the Schmutzi MT contamination analysis) (*55*) since the concordance of higher MT contamination (0.06-0.12) samples had lower concordance according to our experiments using high coverage aDNA data (data not shown).

We used the ancIBD (version 0.5) python libraries with the Python 3.6.8 environment for IBD fragment analysis (*68*). Phased and imputed variants of experimental aDNA samples were post-filtered to include only the positions of the 1240K AADR marker set and lifted to the hdf5 data format as described in the ancIBD manuscript. IBD fragments were identified with the default parameters recommended for aDNA analysis (emission model haploid_gl2, HMM model FiveStateScaled, and the p_col=’variants/RAF’ option to use GLIMPSE2 reference AF data from the imputed variants). We identified IBD fragments >=8cM. The remaining IBD fragments were filtered to include only IBD fragments with >= 220 SNP/cM marker density as recommended in the ancIBD manuscript.

Shared IBD network was generated in R, with the application of packages ggplot2 (https://ggplot2.tidyverse.org/) (*69*), and igraph (https://igraph.org/) (*70*), with Fruchterman-Reingold weight directed algorithm.

## Acknowledgement

We are grateful to Miklós Kásler for supporting the excavation and the genetic analysis. We owe a debt of gratitude to Imre Küzmös for his help in the yfull phylogenetics. We thank Gelegdorj Eregzen for the sample of the elite Xiongnu individual (AG6F). This research was partially funded by the Competence Centre of the Life Sciences Cluster of the Centre of Excellence for Interdisciplinary Research, Development and Innovation of the University of Szeged to T.T., Z.M. and E.N. (the authors are members of the ‘Ancient and modern human genomics competence center’ research group). This research was funded by grants from the National Research, Development and Innovation Office (TUDFO/5157-1/2019-ITM and TKP2020-NKA-23) to E.N. This work was partially supported by ÚNKP-23-4 (grant agreement no. ÚNKP-23-4-SZTE-650) New National Excellence Program of the Ministry for Culture and Innovation from the Source of the National Research, Development and Innovation Fund to B.T.

## Supplementary file

It contains Supplementary text 1-3 and Extended figures 1-4

## Supplementarx texts

**Supplementary text 1**. The archaeological description of Abasár, Bolt-tető site

**Supplementary text 2**. Archaeological description of the studied burials

**Supplementary text 3**. Radiocarbon dating of the remains at Abasár Bolt-tető site

## Extended figures

**Extended figure 1**. Reconstructed equivalent family trees of the larger family from Abasár Bolt-tető site based on the results with correctKin.

**Extended figure 2**. Figure 5 supplemented with sample IDs.

**Extended figure 3**. European PCA of Abasár samples (red squares) supplemented with the individuals from the Báthory cemetery at Pericei site (blue triangles) and the two Báthory family members (green triangles). The overlap between the two cemeteries indicates similar genome pattern for the two medieval Hungarian aristocratic group.

**Extended figure 4**. Eurasian_PCA of Abasár samples supplemented with the Conqueror cline (light blue crosses) and Conqueror Asia Core samples (blue squares). The cluster of Abasár samples overlaps with the Conqueror cline, indicating minor Asian patterns within their genomes feasibly originating from the conquerors.

## Supplementary tables

**Table_S1_Abasár_Stats_Summary**

Table S1a_Archaeological data of the studied samples

Table S1b_Low coverage shotgun data of the samples from Abasár - Bolt-tető.

Table S1c_Whole genome NGS data of the samples from Abasár - Bolt-tető.

Table S1d_List of SNP-s provided against rCRS and SNP used to derive mitochondrial haplogroups using HaploGrep (24).

Table S1e_*Post mortem* demage (PMD) measurement of the sequenced libraries.

**Table_S2_kinship**

Table S2a_Results of kinship analysis with correctKin.

Table S2b_Results of kinship analysis with READ algorithm.

**Table_S3_ Phylogenetic analysis of y Hg N1a1a1a1a4**

**Table_S4_ List of the samples applied in Unsupervised ADMIXTURE analysis**

**Table_S5_PCA_backgrounds**

Table_S5a_List of the background modern samples of the European PCA

Table_S5a_List of the background modern samples of the Eurasian PCA

**Table_S6_qpAdm**

Table_S6a_List of populations included in the qpAdm analysis

Table_S6b_List of the samples from the left populations

Table_S6c_List of the samples from the right populations

Table_S6d. Two-source qpAdm modelling of the Abasár samples

**Table_S7_List of shared IBDs between pairs of samples**.

The threshold of IBD length was determind in 8 cM.

