## Supplementary text for "Archaeogenetic analysis revealed East Eurasian paternal origin to the Aba royal family of Hungary"

### Archaeological and historical background

#### Supplementary text 1. The archaeological description of Abasár, Bolt-tető site

The Abasár, Bolt-tető archaeological site has been known since 1954, when the construction of a fire station on the previously undeveloped inner-city hilltop in Abasár revealed Árpád-era stone walls and tombs by the researchers of the Eger Museum[1,2]. Already at this time, carved column capitals of such quality were unearthed that made it clear that the hilltop is the location of the Benedictine monastery founded by Sámuel Aba in 1042, where eventually the king himself was laid to rest[3,4]. During the communist and socialist eras, several buildings were constructed on the hilltop, unfortunately without archaeological excavation or artifact preservation, despite the clear archaeological significance of the area.

In the 1950s, the residence of the president of the Egerszólát Farmer's Co-operative (922nd lot) was built, followed by the cultural center of the village (958th lot) and Aba Borozó (957th lot) around 1960, all on a 600m2 area. In the '70s, another residential building (953/3rd lot) and a winemaking processing building with three levels below ground (952nd lot) were added. The basement of the president's residence destroyed one-third of the circular rotunda turned out to be an earlier building than the Benedictine monastery, along with the early cemetery around it. The cultural centre and the wine bar caused immeasurable damage to the Benedictine monastery wings, and the other residential building damaged the nave of the monastery church and partially covered it. The winemaking processing building destroyed the cemetery around the church.

In the unbuilt area, archaeological work took place in 1970-71 in connection with the landscaping of the courtyard of the cultural centre. At that time, Nagy Árpád uncovered important sections, but due to the early death of the archaeologist and poorly documented excavation, only limited excavation documentation remains, including a three-page excavation report and some detailed drawings stored in the Hungarian National Museum archive from 1971[5,6,7]. Based on these, it can be seen or inferred that the circular rotunda's almost fully excavatable area was uncovered at that time. Nagy also found the eastern, straight-ended sanctuary wall of the Benedictine monastery church, the walls of the side chapel added to the sanctuary from the south, a spiral staircase, and the presumed wall of the western closure of the monastery church. Numerous graves and three carved tomb covers were also found during the excavation, some of which were located in the middle of the side nave next to the sanctuary, near the spiral staircase. Although Nagy Árpád uncovered significant areas and important findings, the lack of documentation and an overall excavation plan means we don't know exactly where and what he excavated – we can only infer from the current excavation results that he may have excavated in certain areas, but it is certain that he did not dig under the parking lot poured in 1970.

After 1971, the area was landscaped, and despite significant findings, no further archaeological excavations took place. Sámuel Aba's Benedictine monastery did not receive further attention or financial resources at that time. Local residents were dissatisfied with this situation, and in 2006, forming an association, they excavated the rotunda section that was not destroyed by the construction of the president's residence[8]. This work, involving local volunteers and students from Eger, took place intermittently until 2011 when a temporary roof was placed over the excavated area of approximately 100m2[9]. However, the work did not continue, and the protective roof aged, leading to the deterioration of the exposed ruins.

A year after the establishment of the Institute for Hungarian Research, in the fall of 2020, the dismantling of the protective roof and a more comprehensive, interpretable excavation of the site began at the initiative of the local government in 2019. The work continued at full speed in 2021, and parallel to the excavation, efforts began to conserve the uncovered artifacts. In 2021, the president's residence on lot 922 was dismantled, and the remaining, partially destroyed parts of the rotunda were excavated. The years 2022-2023 focused on the conservation of the excavated walls and structures, and by the end of 2023, the floor plan reconstruction of the rotunda was completed, along with the preservation and partial reconstruction of the sanctuary and side chapel of the Benedictine monastery church and one room in the eastern wing of the monastery – making part of the excavation area partially accessible to the public.

**New Archaeological Results**

During the 2020-2021 excavation, our knowledge expanded significantly, as the previously explored area of 100m2 increased to nearly 1800m2. The excavation was conducted by Ásatárs Kft. under the archaeological leadership of the Institute for Hungarian Research led by Miklós Makoldi, with the overall leadership of Zsolt Gallina and Gyöngyi Gulyás. The anthropological material recovered from the site underwent genetic analysis at the Archaeogenetic Research Center of the Magyarságkutató Intézet, led by Endre Neparáczki and Gergely Varga. Here are the main results of the excavation:

**1.1. The Rotunda**

During the excavation, it became clear that the earliest building on the site is the rotunda, partially destroyed by the President of the Agricultural Cooperative. It is fully oriented from East to West and is surrounded by a cemetery consisting mainly of tombs lined with stone slabs, which were partially overbuilt by the 1042-founded Benedictine monastery – the orientation of the monastery buildings differs (northeast-southwest), indicating that the rotunda predates Sámuel Aba's constructions. This also suggests that the rotunda is earlier than Sámuel Aba's constructions, including his tomb. This is supported by the fact that no graves were found inside the rotunda, and the rock surface in the interior was untouched, contradicting the earlier belief that Sámuel Aba was buried here. The dating of the rotunda will likely be determined through isotopic analysis of the early stone-lined graves around it.

**1.2. The Monastery Church and the Monastery**

The most significant discovery in 2020 was the identification of a large, straight-ended sanctuary to the southwest of the circular church. This sanctuary belonged to a late-Romanesque style church, associated with the Benedictine monastery founded by Sámuel Aba in 1042. The excavation revealed three periods of the monastery church, notably the late-Romanesque single-nave church with straight-ended sanctuary (KT2 period), a possible 15th-century floor in the same church (KT3 period), and the original 11th-century remains of a two-towered church with a three-nave, semi-circular sanctuary (KT1 period).

**The Monastery Church's First Period (KT1)**

To the southwest and west of the round rotunda, previously built on the eastern edge of the Bolt Hill rock dome, the Benedictine monastery founded by Sámuel Aba in 1042 was established, intended as the burial place for the king. Originally, the monastery church was built as a three-nave, circular apse, two-towered structure, measuring 10x30 meters, which was almost completely destroyed during the Mongol invasion. During the archaeological excavation, the outer wall plane of the circular apse, "protruding" to the east from under the straight-ended apse of the 13th-century period (KT2 period), and the protruding foundation plane were clearly identified. The walls of the apse of the KT1 period, which facilitated the calculation of the radius, were found during the excavation. Unfortunately, the apse of the side aisles is either completely covered by the 13th-century masonry (KT2 period) or destroyed by burial pits to the rock surface—hence, deductions can be made regarding the semi-circular apse of the side aisles. Additional clues to the early period are the square cross-sectioned Budakalász limestone column at the northeastern corner of the church, originating from the northern aisle of the KT1 period, which was entirely walled into the northern apse wall of the KT2 period. Similarly, a large Budakalász limestone "in situ" block found at the eastern end of the northern wall of the nave (KT1 period) provides a reference point for the early church's northeastern corner.

In the apse of the main nave of the KT1 period, the supporting pillar of the 13th-century period altar stone (KT2 period) is centrally located, suggesting that the early period may have had its altar for celebrating Mass. Among the pillars at the intersections of the aisles of the KT1 period, only the foundations of two remain, as the rest were destroyed by later constructions and later burials. However, at the western end of the church, the massive, high foundations of a pair of towers connected from the inside to the western gable wall of the church were clearly visible, indicating robust towers.

The floor level of the KT1 church could only be sporadically identified just above the rock surface. According to the archaeological excavation, the church floor was likely at a single level and may have been covered with stone or terrazzo, although in most places, only the dusty floor indicated this, with remnants of terrazzo plaster remaining in one or two corners.

Sámuel Aba may have been buried in the main nave of the KT1 church, as this was the holiest place in the Benedictine monastery he founded for burial. In the central line of the main nave, there were several rock-cut graves suitable for the royal burial, but without exception, these contained burials from later centuries, according to C-14 dating. Unfortunately, Sámuel Aba's remains, buried with royal accompaniments, were not found, which is not surprising given that the Mongol devastation almost completely destroyed this church to the floor level, likely disturbing the graves as well. In the geometric center of the KT1 church, a large burial pit was found, with later burials, equipped with 15th-century tomb slabs, indicate, but the size and position of the burial pit could have been suitable for Sámuel Aba's royal burial place, which unfortunately did not survive due to the Mongol destruction.

A monastery wing was also attached to the KT1 church from the north, the remnants of which were found under the walls of the northern side of the church's apse during the 13th-century alterations, with the same floor plan as the 13th-century monastery. These monastery remnants only remained 1-2 courses high compared to the original 11th-century floor, which we found under the Gothic floor levels. However, the excavation successfully identified the remains of the monastery belonging to the KT1 period as well. This monastery, too, was likely a square-shaped building complex with an internal courtyard, similar to its counterpart rebuilt in the 13th century. It is important to note that even this early monastery (KT1 period) was built on top of the earlier cemetery, with graves around the earlier rotunda to the east. Therefore, we believe that the rotunda could be much older than the KT1 period. The further exploration of the monastery wings is greatly hindered, of course, by the building of the cultural house, which partly conceals and partly destroys it and is still standing today.

**The Side Chapel Attached to the KT1 Period Church from the East**

After the completion of the KT1 period early church and the northern side of its apse, a small side chapel was added from the east, which connected to the sanctuary of the northern side aisle of the KT1 church from the east. It became clear during the archaeological excavation that Nagy Árpád likely already explored the chapel, but between 2006 and 2011, the northern half of the building and the tomb inside were definitely excavated since, at the beginning of the excavation by the Institute of Hungarian Research in 2020, this area was open and exposed under the protective roof.

The small building is divided into a straight-ended "sanctuary" and a small nave, with internal buttress-supported vaulting likely holding a cross vault over the sanctuary. In the middle of the nave, there is a grave, which was either excavated by Nagy Árpád or during the excavations of the early 2000s—hence, the bone material cannot be identified, although the person buried there may have been significant.

The building's construction history is clearly visible in the results of the 2020-21 excavation. It is evident that the building was added to the semi-circular apses of the KT1 period church, but it is not contemporary with them. The same can be observed at the square-shaped stone at the northeastern corner of the KT1 church, where it is clearly visible that the corner element was added to the chapel wall from the east. However, it is also apparent that the straight-ended apse wall and the northeastern support pillar of the KT2 period monastery church were already built on top of the chapel wall—thus, the building could no longer stand at that time, and the external ground level was higher after the Mongol devastation than the ruined chapel walls.

From all this, it can be inferred that the chapel may have been built sometime in the second half of the 11th century, but the Mongol invasion of 1241 razed it to the ground, and the building was not reconstructed. Instead, the grand KT2 church was erected on top of the ruins, leaving only the part below the external floor level of the KT2 church preserved. Additionally, it is observable that part of the lintel of the chapel's doorway remains in the western direction—thus, it is certain that the chapel could be accessed through a door created by breaking the apse wall of the northern side aisle of the KT1 church, and its entrance opened from the sanctuary zone of the first-period monastery church. This suggests that the person buried here could be identified with one of the important members of the Aba lineage who died between 1060 and 1240 or possibly with one of the abbots of the monastery. Unfortunately, the bone material recovered from here cannot be identified today due to the lack of documentation.

**The Second Period of the Monastery Church and Monastery (KT2)**

The Benedictine monastery founded by Sámuel Aba underwent its first destruction during the Mongol invasion, as it fell within the main route of the Tatar armies heading towards Buda, resulting in the devastation of both the monastery church and the monastery itself. Interestingly, there is no apparent destruction or reconstruction up to a height of 1 meter 50 centimetres on the walls of the rotunda. Of course, the vault and higher walls of the rotunda could have also been destroyed, as the Mongols surely did not spare this building either. However, the destruction was elemental. The KT1 period church and the monastery were almost completely destroyed.

Nevertheless, after the Mongol invasion, the Aba clan embarked on a gigantic construction project. The monastery church underwent significant alterations and was reconstructed, along with the monastery wings. However, the burial chapel connected to the KT1 church from the east was never rebuilt. The walls of the KT1 church were uniformly dismantled up to a height of 50 centimeters from its internal floor level, and new external and internal floor levels were established, higher than the original.

The KT2 period church evolved into a late Romanesque style, single-nave church with corner pilasters, and the internal dimensions of the sanctuary were 9x18 meters. The length of the church could reach up to 30 meters, but the complete area is not yet excavated, so the exact data is unknown; however, the width of the nave is 12 meters. With these dimensions, the straight-ended pilastered sanctuary, and the large monastery church with divided nave using the western wall of the KT1 church as a dividing wall, the KT2 monastery church shows many parallels in its layout, size, and form with the first period of the Dominican monastery church on Margaret Island, which was built by Béla IV as the monastery and final resting place of his daughter, Saint Margaret. Moreover, later, King István V was also laid to rest in this church[10,11,12]. The size, quality, and similarity of the two churches and monasteries indicate that the Aba clan, not prominently featured in historical sources in the 13th century, had financial resources after the Mongol invasion comparable to those of the king, who, due to the country's escape from the Mongols, built a monastery for his daughter with royal splendour!

The wealth of the Aba clan is also indicated by the painted marble fragment found under the southern side altar of the church sanctuary, depicting the Virgin Mary with the infant Jesus. The quality and beauty of the carving can be compared to the finest late Romanesque to early Gothic European stone carvings, unparalleled in Hungary until now. This stone carving might have been the altarpiece of the XIII. century monastery church (KT2) dedicated to the Virgin Mary, during its reconstruction after the Mongol invasion.

The KT2 period monastery is essentially a completely rebuilt multi-level monastery constructed on the almost ground-level ruins of the KT1 monastery. The monastery has a courtyard with a brick-paved cloister and a well carved into the rock in the centre of the courtyard. The floor plan study may still pose many questions, as only the initiation of the eastern wing of the monastery and a part of the monastery courtyard fell into the excavated 1800m2 area, and the well of the monastery courtyard, accessible from the cellar system below the site, was covered during the construction of the cultural house but fortunately not entirely or only partially walled up.

**The Gothic Chapel Connected to the Sanctuary of the KT2 Church from the South**

After the Mongol invasion, a large, solid, multi-level chapel was added to the southern wall of the sanctuary of the KT2 church, built in early Gothic style. The southern corners of this chapel were supported by diagonal, large-area Gothic pillars. The construction of the chapel presumably took place in the 14th century. During the construction of the chapel, they pierced the southern wall of the sanctuary of the KT2 church, making the ground floor of the chapel accessible. In the centre of this ground floor, a large burial pit is located, which Árpád Nagy already excavated in 1971[13]. Presumably, it was from here that the Gothic tombstone decorated with a 14th-century lace cross, currently inventoried in the Mátra Museum in Gyöngyös in an unidentified state, was unearthed. The carved stone floor chapel must have been two stories high, as the foundation of a stone spiral staircase was found in its southwest corner. Árpád Nagy also mentioned this staircase in his report, describing that a piece of carved staircase element is still in place... Unfortunately, this stone has disappeared over time.

The Gothic two-story burial chapel could have been built for a person of high rank, as it required the disruption of the southern wall of the sanctuary of the KT2 church. The existence of a two-story building indicates a very high prestige. It is assumed that this 14th-century chapel was built for none other than Amádé Aba, one of the most famous members of the Aba clan and a powerful oligarch of the 14th century, possibly before his death. Unfortunately, the identification of the skeletal remains from the ground floor of the chapel seems hopeless due to the lack of documentation.

Perhaps related to the vault of this chapel are two fragments of 14th-century Gothic rib vaults, each with two ribs. These fragments were used to support, or possibly horizontally align, a carved, Aba-coated tombstone with an inscription located north of the main altar of the KT3 church. This repositioning and levelling took place after the Hussite destruction in the second half of the 15th century.

**The Third Period of the Monastery Church and Monastery (KT3)**

In the life of the monastery, the second period of destruction and subsequent reconstruction occurs in the 15th century under circumstances that are not exactly clear. In any case, the major openings, especially the doors of the monastery, are replaced with finely carved eyebrowed stone-framed doors, which, in several cases, are substituted for the original 13th-century openings, sometimes altering their size. A good example of this is the doorway leading from the inner courtyard of the monastery to the rotunda, used as a baptistery at the time, where the wide stone threshold is significantly narrowed and fitted with a smaller-sized, fine-arched Gothic stone frame in the 15th century, with the original threshold left in place.

During this time, the interior of the monastery church also receives a new brick covering, laid with 20x20x5cm Gothic floor tiles.

But what could have been the reason for the 15th-century renovation? Most likely, a serious devastation, as evidenced by several factors within the church and the monastery. The most striking evidence of serious devastation and the accompanying massacre is found in the upper layers of the 15th-century carved stone-covered graves in the sanctuary of the KT2 church. Masses of scattered human bones are discovered, and often partial human remains are found in anatomical order, indicating that after the construction of the graves, a complete destruction ensued on the monastery grounds. It seems that everyone was slaughtered to such an extent that there was no one left to bury the dead. The corpses could have lain unburied for months, partially disintegrated by wild animals. Later, those returning to the monastery threw these human remains into the graves with hinged stone covers, in some cases as scattered bone remains, and in other cases, the remains of bodies somewhat preserved in anatomical order.

Such devastation in the 15th century could have only been carried out by the Hussites who repeatedly invaded the area and caused significant damage to the fortress in Kisnána. Aba's relative, Péter Kompolti, already falls in battle against the Hussites in 1420[14,15,16]. The incursions likely continued until the time of King Matthias[17]. It is probable that the benedictine monastery in Abasár was destroyed in this period and was probably rebuilt during the era of the Hunyadis, where renovations took place, such as the retiling of the monastery church with bricks or the replacement of door openings in the monastery wings with Gothic stone-framed openings.

An indication of the use of Gothic door frames on the upper floor of the monastery is an uncovered, scorched door frame among burned wooden beams in the northern wing of the monastery. This suggests that there might have been Gothic door frames on the upper floor of the monastery since, in the vicinity of the fallen frame on the ground floor, has no wall opening.

It is evident that the Aba clan still had significant financial resources in the 15th century to rebuild the church and monastery, incorporating openings that matched the style of the time. This is not surprising, given that the Nánai branch of the Abas, who were probably donors to the monastery at this time, are frequently mentioned in royal positions in contemporary sources[18,19,20].

**The Destruction of the Monastery**

After the 15th century, there is scarce historical information about the life of the monastery or the Nánai Kompolti family who owned it. However, archaeological findings indicate that life in the complex did not cease. According to numismatic evidence, the monastery was used until the 1630s when, in all likelihood, a Turkish invasion put an end to its existence.

What is certain is that the devastation during the Turkish era involved intense fires, and even the Gothic door frames were found severely burnt and collapsed on-site. After the Mongol and Hussite devastations, the church and monastery were rebuilt, but they succumbed to the Turkish invasion. The area remained uninhabited and undeveloped until the 20th century. The ruins were likely visible until the 19th or early 20th century.

In the 17th century, traces suggesting bronze casting and blacksmith activities were found among the ruins. Subsequently, there were wine-making developments and constructions associated with the active use of the underground cellar system beneath the ruins. Large wine cellars and pressing houses were built in the field of the ruins.

However, the development and destruction of the site only began in the mid-20th century with the construction of József Dér's house, followed by the main building of the cultural centre. These constructions and subsequent gradual urbanization posed significant challenges in excavating, interpreting, and presenting the ruins.

8 Dobó István Vármúzeum Régészeti Adattára: RA. 1212 Ásatási jelentés Fodor László ásatásáról.

9 Dobó István Vármúzeum Régészeti Adattára: Rad. 1563-2013. Ásatási jelentés Fodor László – Lovász Emese ásatásáról

10 Almási Tibor (2000): A tizenharmadik század története. Budapest: Pannonica Kiadó.

11 Kristó Gyula (2007): Magyarország története 895-1301. Budapest: Osiris Kiadó.

12 Attila Zsoldos: The Árpáds and Their People. An Introduction to the History of Hungary from ca. 900 to 1301. Budapest: Research Centre for the Humanities

13 Dobó István Vármúzeum Régészeti Adattára: RA 358. Ezen a helyen Nagy Árpád megemlíti, hogy a mellékkápolnában két sírlappal fedett és éremmel datált sírt is feltártak A XIII. század elejéről és a XIV. századból.

14 Buzás Gergely (2013): A kisnánai vár története. Visegrád, Magyar Nemzeti Múzeum Mátyás Király Múzeum.

15 Buzás Gergely - Kovács Olivér (2016): Középkori várak. Budapest, Magyar Nemzeti Múzeum.

16 Nováki Gyula - Baráz Csaba - Dénes József - Feld István - Sárközy Sebestyén (2009): Heves megye várai az őskortól a kuruc korig. [Magyarország várainak topográfiája 2.]. Budapest-Eger: Castrum Bene Egyesület - Bükki Nemzeti Park Igazgatóság.

17 Egy 1460-as adat szerint a husziták ekkor foglalják el Gyöngyöspata települést. (Szabó János Győző: A gyöngyöspatai Szent Péter templom. In: Az Egri Múzeum Évkönyve - Annales Musei Agriensis 21, 1985. 9. Az eredeti hivatkozott oklevél a Magyar Nemzeti Levéltárban található OL DF 250487 számon.)

18 Buzás Gergely (2013): A kisnánai vár története. Visegrád, Magyar Nemzeti Múzeum Mátyás Király Múzeum.

19 Buzás Gergely - Kovács Olivér (2016): Középkori várak. Budapest, Magyar Nemzeti Múzeum.

20 Nováki Gyula - Baráz Csaba - Dénes József - Feld István - Sárközy Sebestyén (2009): Heves megye várai az őskortól a kuruc korig. [Magyarország várainak topográfiája 2.]. Budapest-Eger: Castrum Bene Egyesület - Bükki Nemzeti Park Igazgatóság.

**Supplementary text 2. Archaeological description of the studied burials**

During the excavation at Abasár Bolt-tető site, more than 300 burials were uncovered within and outside of the gothic church building (Object 7). As the indoor graves potentially harbour aristocratic individuals' skeletons, we took those into archaeogenetic investigation to identify the potential Aba clan members. Below we present the archaeological description of the studied indoor burials. Since various construction phases of the temple were identified during the excavation, the archaeological phenomena uncovered in each layer were separated with stratigraphic numbers (Snr). We start with the description of the most important burials, whose findings were genetically characterized in the main text.

**1)** The prominent burial in the sanctuary with the carved and scripted cover stone, labelled with blue quadrate in Figure 1. This includes Snr 8, Snr 57 and Snr 58/1-2.

**2)** The two graves in the sanctuary labelled with yellow quadrate in Figure 1. This includes Snr 55/a-b and Snr 59/a-b.

**3)** The double grave in the geometrical centre of the church labelled with green quadrate in Figure 1. This includes Snr 194/a-d, Snr 261 and Snr 262.

**4)** All other graves within the temple. These include Snr 200, Snr 201, Snr 309, Snr 310, Snr 311/a-b, Snr 316, Snr 341, Snr 390, Snr 401, Snr 445, and Snr 450.

**1.1) Snr 8**

Archaeological dating: 15th century CE

Burial with a tombstone in the temple (Figure S1), towards the east from the stone stairway and towards the northwest from the altar foundation. At the northeast end, a brick floor is observed. In the northwest section, there is a stone frame (straight-moulded, Gothic vault),

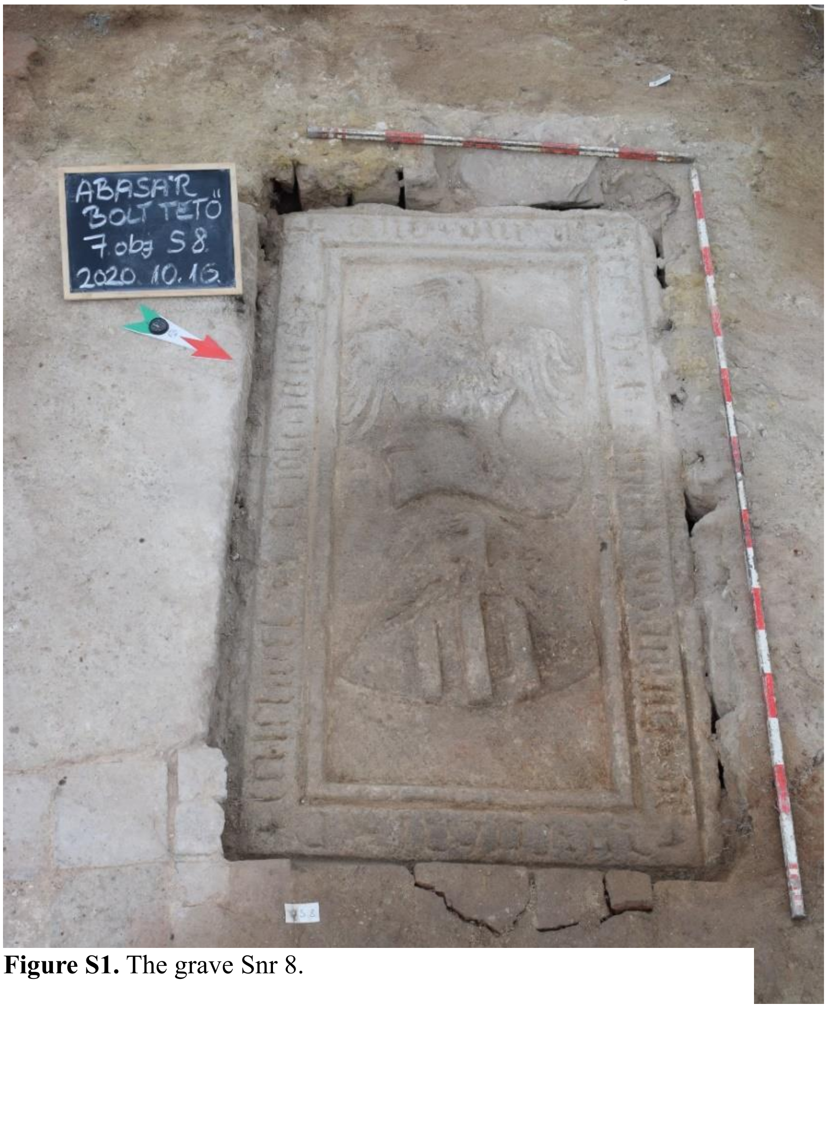

**Figure S1.** The grave Snr 8.

squared stones to the southwest, a larger stone slab to the southeast, supported by squared stones, likely added later. The cover stone is 190 cm long, 100 cm wide, and 25 cm thick. The original cover stone displayed the crest of the Aba clan and a presumably readable Gothic minuscule inscription in a circular pattern: 'In the year of the Lord MCCC?, here rest János and ………… sons: Mihály and János.' Unfortunately, the last numeral of date inscribed with Latin numbers was damaged and could not be decode, thus, the date can be equivalently translated as 1401 or 1405 or 1450. Based on the year and names, we can likely assume these are the sons of János, who was a member of the Aba clan.

The cover stone was lifted with a machine, revealing a layer of 50 cm thick stone rubble underneath. From the level of appearance, down to a depth of 60-70 cm, five jumbled human skulls lay next to the northwest wall. Unfortunately, the bones in the grave were disturbed, but at the bottom of the grave, we found two relatively intact skeletons in anatomical positions (Snr 56, Snr 57). As these two remains could be unambiguously regarded as primary burials, based on the preliminary archaeological data it is assumable that they were Mihály and János. Above these skeletons, however, we found the mixed bones of more than a dozen individuals and a few anatomically ordered body parts, suggesting that these bones may be the remains of untended dead collected after some larger massacre or the exhumed bones of earlier burials (perhaps relatives of the Kompolti branch brought here during the devastation of the Kisnána Castle in the Hussite attack of 1470?). After extracting the Snr 57 skeleton, we found human remains likely placed in a chest or coffin at the same time and then buried (Snr 58). Under Snr 58, the stone floor of the burial chamber, breaking in the middle, began to deepen in a square area of approximately 90 x 110 cm. Here, at a depth of about 140 cm, we found human bones throughout the depression. The depression may be an early ossuary (Snr 60), possibly contemporaneous with the early period of the temple, containing the scattered bones of several dozen individuals – along with a fragment of spiral-lined early Árpádian pottery.

From the grave, iron nails, a glass bead, a coin, an 'L'-shaped right-angled iron clasp, round stained glass(?) inserts, and glass melts were unearthed. Fabric remnants were found in the southwest corner of the grave. In the south section, many iron nails, 'L'-shaped iron clasp with round lead inserts, marble pieces, and glass melts were found. From the filling of the grave, a coin from the western corner emitted by Matthias (1463) also emerged. Carved stone finding: Gothic arch – vaulting element.

Sampled remains from the Snr8: HUAS81, HUAS82, HUAS83, HUAS84, HUAS85, HUAS86, HUAS87, HUAS88, HUAS89 and HUAS57F

**1.2) Snr 57**

As indicated above, this is a sub-feature of Snr 8, with archaeological dating: 15-16th century CE.

In the grave marked Snr 8 (on its north side), bones were found approximately 90 cm deep from the level of the grave's appearance. Adjacent to it was the skeleton marked Snr 56 (Figure S2). Beneath it, bones belonging to several individuals from Snr 58 were uncovered (HUAS58.1 and HUAS58.2). The burial of the individual in Snr 57 (HUAS57) was lying on its back in an extended position. Only a small piece of the skull remained, with the left forearm bones missing. The right lower leg was positioned among the femurs of Snr 56. The bones of the upper body were disturbed. Orientation of Snr 57 (HUAS57): Southwest to Northeast. No grave goods were present. As it was already mentioned in the description of Snr 8: it is conceivable that at the bottom of the grave, two relatively intact skeletons lying in anatomical positions can be identified as János's sons, Mihály and János. One of them, marked Snr 56 was not sampled, because of the lack of skull.

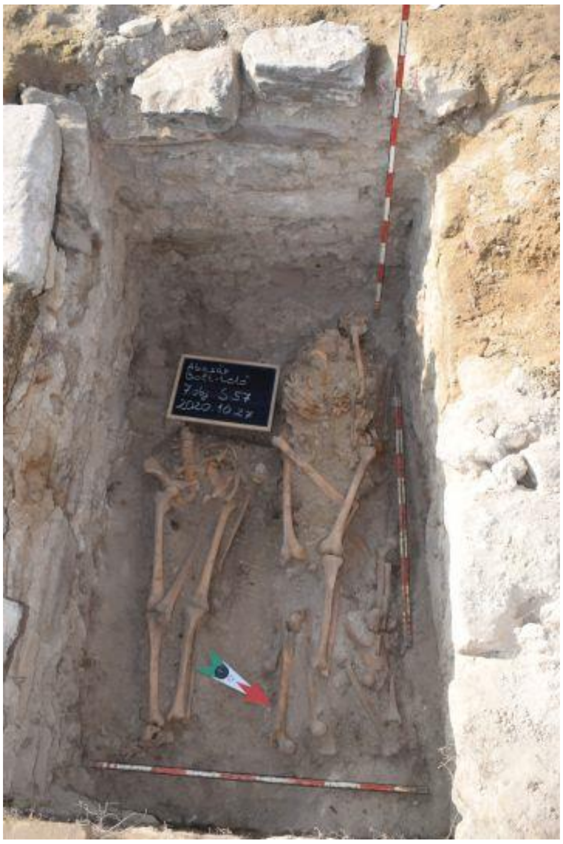

**Figure S2**. The skeletons of Snr 56 (left) and Snr 57 (right).

#### 1.3) Snr 58/1-2

Archaeological dating: 15-16$^{th}$ century CE. Graves or ossuary.

In the temple, in the grave marked Snr 8 (on its north side), various skeletal parts belonging to multiple individuals were uncovered beneath the Snr 57 skeleton. These skeletal parts were contiguous but not in anatomical order. Since the bones appeared on a regular rectangular surface, it is conceivable that they were gathered in a coffin and placed in the pit (Fig. S3). Several femurs (belonging to at least 3 individuals), forearm bones, pelvic bones, sacral bones, and smaller bones (vertebrae, ribs) can be mentioned. Studied samples: HUAS581 and HUAS582.

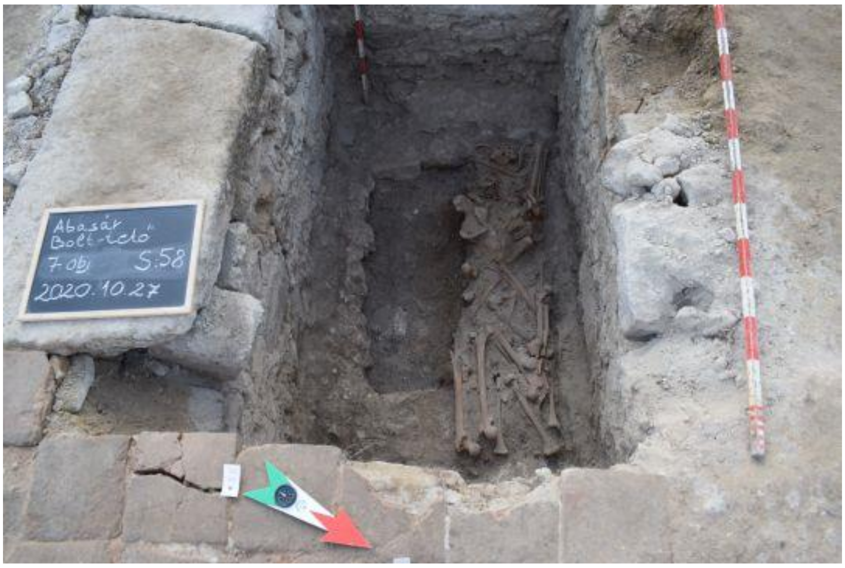

**Figure S3**. The skeletons of Snr 58.

**2.1) Snr 55/a-b**

Archaeological dating: 15-16th century CE

In the temple, a rectangular grave Snr 55 appeared at the corner of the walls. Parallel to it, towards the northwest, lay the burial marked Snr 59 (Figure S4). During the excavation of the grave, the brick floor and the mortar layer underneath were damaged. The appearance level of the grave was sunken, and around it, the mortar layer in which the former brick floor was embedded is clearly visible. Irregular rounded stones were found in the middle and northeast part of the grave. The Snr 55 grave extended into the bottom of the later church.

The burial chamber of Snr 55, constructed with smaller and larger stones in five rows, was revealed. Its bottom was 85-90 cm below the appearance level. The grave pit was a regular rectangle, with steep walls, and the bottom was on the raw rock surface. In the grave, the well-preserved skeleton (HUAS55A) of an adult individual lying on its back in an extended position was found. The skeleton was preserved up to the knees; the lower legs were missing, likely removed during later disturbance. The burial was evident in the eastern wall of the grave. It was a coffin burial, with the coffin visible 50 cm below the appearance level of the grave. Measured dimensions: height 120 cm (up to the knees), shoulder width 37 cm. The facial part of the skull of the Snr 55/a skeleton was damaged, both arms were extended. The left forearm was placed on the left pelvis, and the right forearm on the outer side of the right pelvis. On the left side of the skull and the shoulder area, one of the femurs of the Snr 55/b skeleton was found. Orientation of Snr 55a Southwest to Northeast.

On the deceased's skull and behind it, at the western end of the grave pit, about 10-15 cm higher, human bones belonging to a separate individual Snr 55/b (HUAS55B) were gathered. Among them: a skull, pelvic bones, lower legs, clavicle.

Finds: Árpád-era coin, iron elements of the coffin, more than 10 iron coffin nails, a large quantity of 14-16th-century ceramics, few animal bones. During the excavation, a gilded bronze object was also uncovered, which could be a shepherd's staff or, more likely, the decorated lower part of a goblet. The four protruding, round-shaped parts were decorated with enamel, presumably showing the four apostles. Sampled remains: HUAS55A, HUAS55B

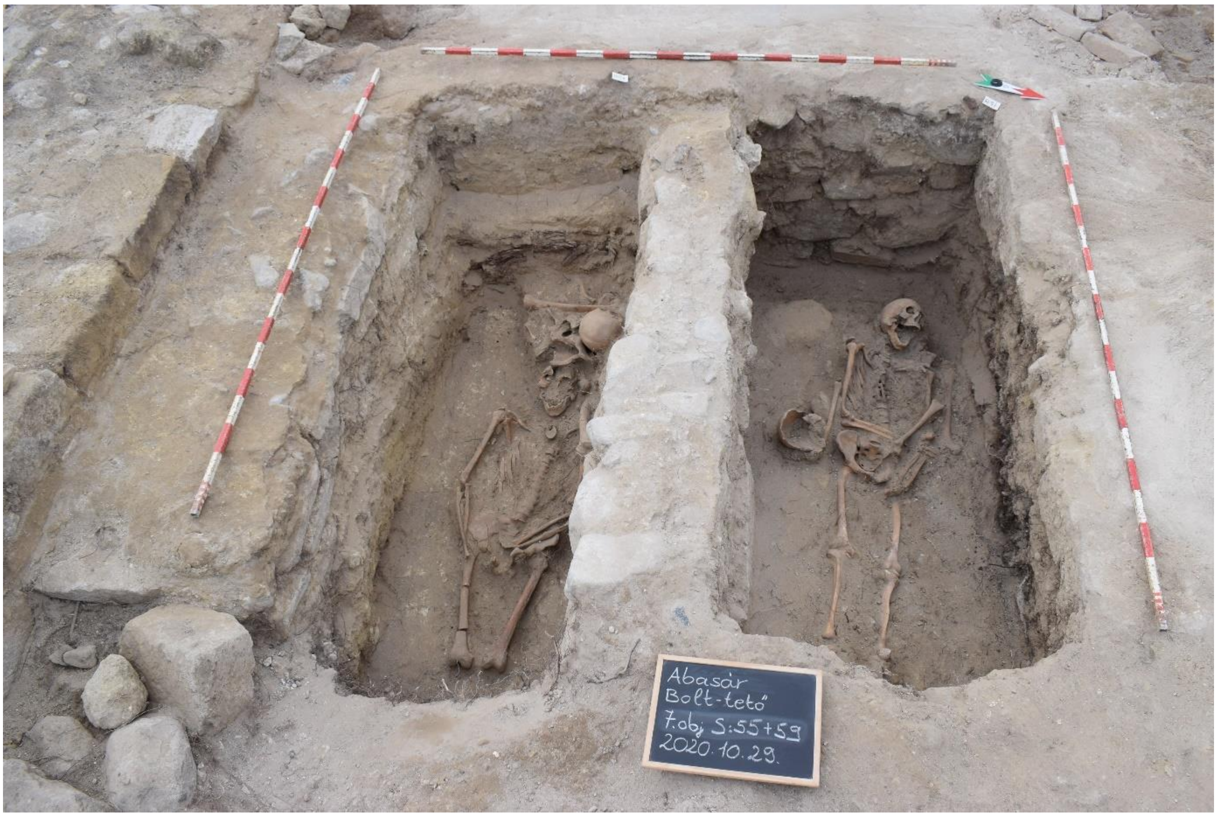

**Figure S4.** The burials Snr 55 (left) and Snr 59 (right).

**2.2) Snr 59/a-b**

Archaeological dating: 14-15$^{th}$ century CE.

In the temple, adjacent to Snr 55, approximately 30 cm to the North, there is a rectangular grave pit (Figure S4). The walls of the burial chamber were crafted from stones, arranged in five rows. In the middle of the grave pit, there lies the skeleton of an adult individual lying on its back in an extended position (HUAS59A), slightly leaning to the left and sunk into the thicker fill beneath. The skull is turned to the left, with the lower jaw fallen off. The right arm is positioned across the hip, and the left hand is placed on the left femur. The grave remains undisturbed. In the vicinity of the bones of HUAS59A the bones of a separate individual Snr 59/b (HUAS59B) were found, with postcranial bones and skullcap. The length of the skeleton is 170 cm, with an estimated height of 165 cm. Shoulder width: 40 cm (38 cm at the elbows). The Snr 59/a skeleton is in anatomical order. Orientation: Southwest to Northeast. No grave goods were present.

Finds: coffin iron bands, few animal bones, iron nails, and a small amount of 14-15th-century ceramics. Samples_HUAS59A and HUAS59B.

**3.1) Snr 194/a-d**

Archaeological dating: medieval. Ossuary

In the geometrical centre of the church a double grave with cover stones with bas-relief decoration depicting the coat of arms of the Aba clan were disclosed (Figure S5). Beneath the cover stones, there is an ossuary (Snr 194). To the north of Snr 194, an eroded layer runs. The human bones were carefully excavated to a depth of 15-20 cm, covering an area of approximately 1.5 x 1 meter. The bones were located in a western to eastern strip. In the eastern part, there were 4 adult skulls (HUAS194A, HUAS194B, HUAS194C, and HUAS194D), and in the central and western part, long bones and other skeletal parts (ribs, vertebrae) were found. Among the bones, there was also an iron nail. All skeletal parts were in secondary positions. During the excavation of the bones, traces of burning were visible on some pieces, which were separately packaged. Below the upper layer of bones, a spinal column and a left pelvic bone were found in situ. During the deepening, beneath the preserved spinal column, there was a bronze casting mould, iron nails, numerous human bones (vertebrae, phalanges, ribs), and 4-5 fragments of medieval vessels.

Approximately 1 meter below the later church's brick floor layer, we excavated another bone deposit in the central part of the cavity. In the southeastern part, there were lower leg bones, pelvic bones, ribs, and one or two vertebrae. On the eastern side, at a depth of 40-50 cm, fragments of a spur were found on the cavity's sidewall. Further pieces, including more fragments of the spur or another spur, were unearthed from deeper layers along with a coffin nail. Additionally, 10-12 iron nails were found, with traces of wood remnants, likely coffin nails. From this depth, only a few pottery fragments (e.g., lid knob) emerged.

Below the layer beneath Snr 194, 15-20 cm deeper, near the southeastern corner of the pit but still 40-50 cm above the Snr 261 grave, an ornate two-part silver belt buckle was found embedded in the wall. Two burials were beneath the Snr 194 ossuary: Snr 261 and Snr 262.

Finds: Above the upper bone layer of Snr 194, there were wheel-thrown, white, thin-walled pottery fragments, iron nails, thin bronze sheets, an iron chest ornament, and wood remnants. Samples: HUAS194A, HUAS194B, HUAS194C and HUAS194D.

**3.2) Snr 261**

Archaeological dating: medieval

The grave Snr 261 was discovered below the ossuary Snr 194. The burial spot was found at a depth of 90-100 cm. Below the bones of Snr 194, a dark brown organic filling is observed.

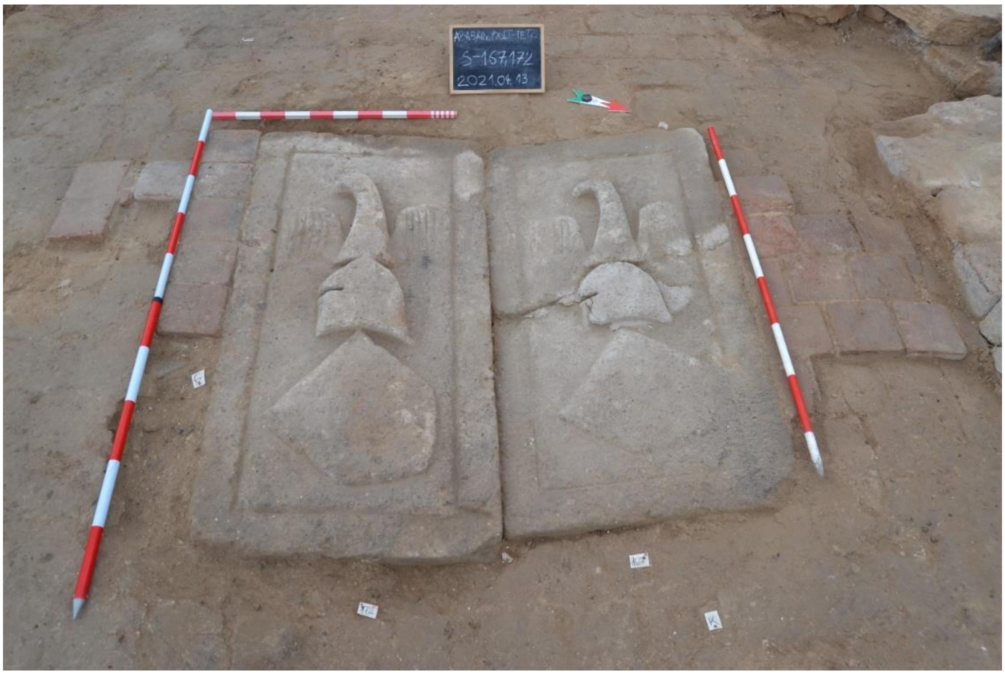

**Figure S5.** The cover stones above Snr 194, Snr 261 and Snr 262.

The edges of the wooden coffin are well-defined, measuring 65-70 x 165 cm. The coffin is wide, and its walls are clearly visible (Snr 261). In the coffin grave Snr 261, there is the well-preserved skeleton of an adult male lying on his back in an extended position (HUAS261, Figure S6). The skull is in its original position, facing upward. The arms are stretched alongside the body. The right forearm is slightly positioned abnormally, as if twisted. The lower leg bones are slightly tilted to the north. Both feet are turned inward. Strong muscle attachment surfaces are visible on the skeleton. As the skeleton was at the bottom of the grave in primary position, presumably this could be the original burial of the grave and the cover stone with the coat of arms of the Abas belonged to this remain.

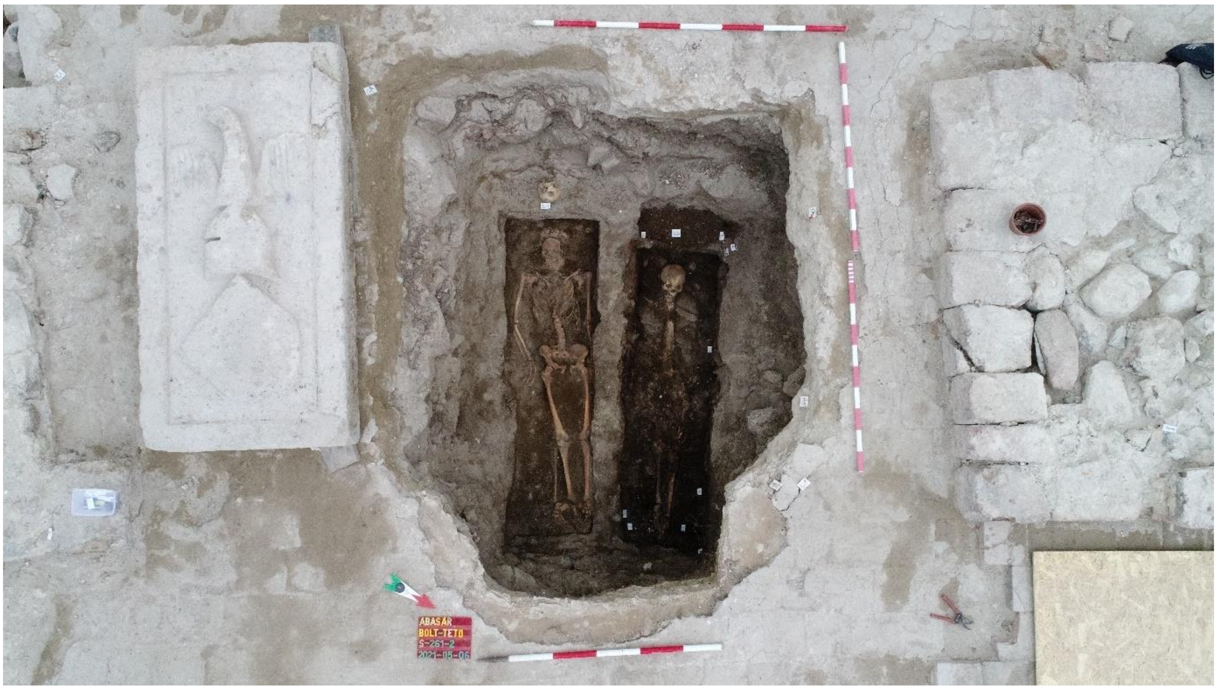

**Figure S6.** The remains of Snr 261 (right) and Snr 262 (left).

Orientation: west to east. No grave goods were present. The coffin nails are preserved. These were positioned on all four sides at equal intervals. The upper nails were at the level of the skull, lying horizontally with the nail heads facing outward. Two nails and an approximately 5 cm long stone wedge were removed during the excavation. At the middle of the northern grave pit wall between Snr 261 and Snr 262, an 'L'-shaped iron reinforcement was extracted. It was secured with approximately 15 iron nails to the coffin. Coffin length: 190 cm, width: 55 cm. Skeleton length: 183 cm. Sample: HUAS261.

#### 3.3) Snr 262

Immediately east of Snr 261, the Snr 262 grave was located (Figure S6). Archaeological dating: medieval

This is the second grave beneath the bas-relief decorated cover stone with the coat of arms of the Abas, which was discovered below the ossuary Snr 194. The burial spot was found at a depth of 90-100 cm with remnants of a coffin. The eastern end of the coffin is brown with organic filling. In the western part, a 3.5 cm wide and 10.5 cm long iron plate was found. In the coffin grave Snr 262, there is the skeleton of an adult woman lying on her back in an extended position with moderate preservation (HUAS262). Smaller stones were laid on the chest and pelvic area. The wall/side of the coffin was well-preserved on the southern side. On the northern side, only the upper remnants, about 5-7 cm thick, were preserved. The skull faced forward, and the arms were not visible due to the collapsed stones and remnants of the coffin. The skeleton was at the bottom of the grave in primary anatomical position suggesting that this could be the original burial of the grave and the cover stone with the coat of arms of the Aba clan belonged to this individual.

During the excavation, two 'L'-shaped iron straps were found, one in the northwest corner and the other in the northeast corner – coffin fittings. Both straps were pierced with 1.5-2 cm long iron nails. Displaced coffin components: 4 iron plates, two of which were bent in an 'L' shape and pierced with iron nails. 3 larger-sized iron nails. Additional coffin components:

Iron strap in the northwest corner of the grave – secured the upper part of the coffin.

Iron nail in the southwest corner – secured the upper part of the coffin.

Iron strap with wood remnants on the northern coffin wall, at approximately elbow height.

Vertically standing iron nail and iron strap at the bones of the right lower leg in the grave.

Sturdy 'L'-shaped iron strap at the northeastern corner of the pit – secured the lower part of the coffin.

Sturdy 'L'-shaped iron strap at the southeastern corner of the pit – secured the coffin. The distance between the lower parts of the 5th and 6th straps was approximately 15 cm, with the 11th finding between them.

Vertically positioned iron nail to the west, about 10 cm from the 6th finding.

Vertically positioned iron nail to the west, about 2 cm from the 1st strap.

Iron strap above the 5th strap – secured the upper part of the coffin.

Vertically positioned iron nail to the southwest of the coffin, about 10-15 cm from the 2nd finding.

Vertically positioned iron nail in the middle of the eastern end of the coffin, showing a large wood remnant.

From the excavation, a wheel-thrown pottery bottom fragment was found. Orientation: West to East. No grave goods were present. Coffin length: 180 cm, width: 50-57 cm. Skeleton length: 165 cm. Sample: HUAS262.

**4.1) Snr 200**

Archaeological dating: medieval

In the vaulted chamber of Snr 192, a grave lies parallel to its northern wall. The Snr 200 skeleton (HUAS200) is mostly *in situ*, with the upper body and skull in their original positions. Other skeletal parts were found in secondary positions. The skeleton likely belongs to an elderly male in a well-preserved state, lying on its back in an extended position. The skull and the chest bones are intact, but the arms are missing. Beneath the skull, another individual's skull rested. Almost directly under Snr 200, there is another skeleton. Stones were placed on the lower extremities, and after removing them, the lower leg bones were revealed. Beneath them, the legs of the deceased underneath were also present. The Snr 200 skull shows sharp, deep incisions with healed edges, indicating that the individual survived the injury.

Orientation: West to East. Grave goods:

On the left side of the chest, near the vertebra, a bronze Parisian clasp.

During the excavation and deepening of Snr 192, 5 small glass beads were found, possibly associated with this skeleton.

Sampled remain: HUAS200.

**4.2) Snr 201**

Archaeological dating: medieval

In the vaulted chamber of Snr192, there is a grave lying parallel to its southern wall (Snr 201). It is situated directly north of the Snr 200 skeleton, potentially contemporary with it. The undisturbed skeleton of an adult male (HUAS201) was lying on its back in an extended position. The skull is intact, and the vertebral column and right ribs are in place, reaching approximately to the sacrum. The arms are missing. After the removal of debris stones at the lower legs (2 carved stones), the bones of the lower legs were exposed. Beside these bones, we uncovered the lower leg bones of at least two more individuals.

Orientation: West to East. Grave goods: none.

Sample: HUAS201.

**4.3) Snr 309**

Archaeological dating: medieval

Near the staircase of the Snr 192 vaulted chamber, to the west, at the same level as the stone level, a grave pit was identified. In the western part of Snr 309, at a depth of 10-15 cm, some human bones (skull, femurs, bones of the upper extremity, and others) were found. Under the stone blocks, a complete skeleton was discovered during the excavation of Snr 309. From the excavation of Snr 309, brick-coloured ceramic and fired, 14th-century pottery shards were found. Under the disturbed bones, additional bones were discovered in secondary position. The Snr 309 skeleton (HUAS309) was located approximately 60-70 cm below the pit's entry level. The pit walls and bottom are well-defined, and it was dug into the raw rock surface. The grave contained the well-preserved skeleton of an adult individual in a supine position. The skull is intact and slightly turned to the right. The left arm is slightly bent, while the right arm is bent at a nearly right angle at the elbow, resting on the pelvis.

Orientation: West to East. Grave goods: 1. Approximately 10 cm above the left lower leg, an iron coffin nail was found, oriented in a North-South direction, with the head to the North. During the excavation, additional 7 nails were uncovered.

Studied sample: HUAS309.

**4.4) Snr 310**

Archaeological dating: medieval

Near the staircase of the Snr 192 barrel-vaulted crypt, to the West from the Snr 309 burial, at the same level as the stone level, a grave pit was identified. The walls of the grave pit were well-defined on the northern and western sides. The fill of Snr 309 and Snr 310 was identical. During the excavation of Snr 310, few human bones in secondary position were found (pelvic bones, ribs, arm bones). In addition, a few nails and a robust metal object were uncovered. In the grave, the well-preserved skeleton of an adult male (?) was found in a supine position (HUAS310). The skull is slightly tilted to the left, and the right side was damaged during the excavation. Both arms were bent at the elbows and placed across the pelvis. The legs were extended, and the feet reached beneath the debris in front of the vaulted crypt.

Orientation: West to East. Grave goods: 1. A bronze coin near the left femur. 2. A bronze coin beneath the left elbow.

Studied remain: HUAS310.

**4.5) Snr 311/a-b**

Archaeological dating: medieval

Directly to the south of Snr 310, within the small stone layer, an irregular rectangular pit was discovered. Its outline was visible at a higher level, and the mortar layer had sunk. Initially, it was excavated in a cross-section, and the western half was exposed. The fill consisted of a 15-20 cm thick layer below the mortar layer, which was less rubble-like and included mortar. Below this, there was a dark brown, rocky-rubble, loose layer. In the grave pit, a few disturbed bones (ribs, vertebrae, bones of the upper extremity) were present (HUAS311B). In the south-western corner of the pit, the skull of an adult individual was left *in situ* (HUAS311A). At the bottom of the pit, the well-preserved skeleton of an adult male in a supine position was found. The humeral bones displayed strong muscle attachment surfaces, indicating robust musculature. The skull was turned to the right, both arms were bent at the elbows, laying on the abdomen. The left pelvic bone and the lower vertebrae had moved out of position. The chest area was depressed, with the skull being the highest point. It was a coffin burial, and the southern part of the pit contained wood remains and iron nails.

Orientation: West to East. Grave goods: 1. From the excavation, two fragments of a bronze coin were found on the outer side of the left lower leg bones, at the same level as the bone. 2. 5 iron coffin nails. Additionally, ceramic fragments dated to the Late Árpádian Age were discovered.

Samples: HUAS311A and HUAS311B.

**4.6) Snr 316**

Archaeological dating: medieval

In the 7th object/7th sector, beneath the mortar layer, a grave was identified. The pit wall was not discernible due to collapsed stones. Human bones were found approximately 70-80 cm below the initial level of the pit. Within the grave, the skeleton of an adult male was found in a supine position with poor preservation (HUAS316). The vertebrae had collapsed, and the skull was absent, with only a small part of the lower jaw remaining. The left upper arm had slightly shifted, unrelated to the excavation process. The right arm was bent at the elbow, lying across the abdomen. The left arm was also bent at the elbow. Smaller fragments of the skull were found among the lower leg bones. After the removal of the skeleton, beneath the left forearm bones and the location of the skull, in the western end of the pit, a dark brown, larger textile remnant was discovered (possible clothing). The largest piece was beneath the skull, measuring 2-3 x 5 cm.

Orientation: West to East. Grave goods: between the feet, a vertically positioned iron coffin nail was found. During the excavation, a small bronze coin was discovered approximately 10 cm above the left upper arm. Near the right ankle bone, a small-sized iron nail was found (possibly a coffin nail).

Sample: HUAS316.

**4.7) Snr 341**

Archaeological dating: Árpádian age

In the Snr 203 grave, skeletal remains belonging to multiple individuals were found. The initial level of the pit was 60-80 cm below, and the Snr 341 skeleton was found in proximity (HUAS341). The skull was positioned higher, with the lower jaw turned down. The facial bones were fractured. Both arms were bent at the elbows, placed on the abdomen. The finger bones were located above the sacrum. The feet were missing.

Orientation: West to East. Grave goods: none. Length of the skeleton: 156 cm (up to the end of the lower leg).

Studied remain: HUAS341.

**4.8) Snr 390**

Archaeological dating: Árpádian age

In the 7th object/7th sector, a grave located West of the stairs was identified. The grave has a rounded rectangular shape with a straight bottom. At the foot end, there were remnants of a decayed coffin. The skeleton found in the grave belonged to an adult individual, lying in an extended position, with good bone preservation (HUAS390). The skull was turned to the left, the arms and legs were extended, and the hands were placed on the neck of the femurs.

Orientation: West-Southwest to East-Northeast. Grave goods: none. Stray findings from the surrounding area: an Árpádian Age pottery shard and an iron band were discovered during the excavation.

Sample: HUAS390.

**4.9) Snr 401**

Archaeological dating: medieval

In the 7th object, beneath the northern side altar, East-Southeast from a wall segment, a grave was uncovered. The Western end was excavated, revealing remnants of a coffin. The remains of at least two individuals were found in the grave. During the excavation, iron nails and pottery fragments from the 14th-15th centuries were discovered.

Orientation: West to East. Grave goods: none.

Sampled remains: HUAS401 and HUAS401P

**4.10) Snr 445**

Archaeological dating: Árpádian Age/medieval

The partial skeleton of an adult individual in the western corner of the Snr 203 grave (HUAS445), located in the 7-8th sector of the temple. The remains were found after the skeletons belonging to other individuals (Snr 354 and Snr 389) were removed from the grave. The preserved part of the skeleton was found on a roughly 40 cm high platform, resting on the raw rock surface at the Western end of the grave. The preserved elements include the skull (with the lower jaw turned outward), 4-6 vertebrae in anatomical order, while other bones were removed during excavation. The dentition is incomplete, with some tooth sockets *antemortem* missing. The skull also displays deformities.

Orientation: West to East. Grave goods: none. Grave pit width: 50 cm.

Studied sample: HUAS445.

**4.11) Snr 450**

Archaeological dating: medieval

The burial in the temple, 7-8th sector, towards the East to Southeast from the Snr 311 grave, a grave pit with a sloping, occasionally uneven bottom, reaching down to the raw rock surface was excavated. The grave contained the remains of an adult male in a supine position (HUAS450), with moderate bone preservation, and occasionally fragmented bones. The skull is compressed and slightly tilted to the left, and the lower jaw has detached. Both arms are bent at the elbows, with hands resting on the pelvis. The legs are extended and moderately preserved, with the feet somewhat turned toward each other at a higher level.

Orientation: West to East. Grave goods: 1. a coin fragment found during excavation (exact location unknown). 2. several iron coffin nails (6-7 pieces). The coffin nails were retained *in situ* from the knee down. Additional nails were found among the ribs and the feet during bone retrieval. A total of 25-30 nails secured the coffin. 3. an iron strap on the outer side of the left upper arm. 4. during skull retrieval, remnants of a copper-alloy fitting were found under the right side. Additionally, an Árpádian Age pottery shard was uncovered during excavation. Grave pit length: 195 cm, diameter: 78 cm, skeletal length: 174 cm.

Sample: HUAS450.

**Supplementary text 3. Radiocarbon dating of the remains at Abasár Bolt-tető site**

Supplementary table 1: Direct AMS radiocarbon dates from Individuals published in this study

| Lab ID | Site | Grave number | Type of sample | Uncalibrated BP | ± | OxCal[1] 95% cal (CE) | Measuring Laboratory[2] | AMS Lab ID |
|---|---|---|---|---|---|---|---|---|
| HUAS55B | Abasár-Bolt-tető | S55B | rib fragment | 696 | 12 | 1276-1300 (92,6%); 1372-1377 (2,8%) | INR Debrecen | DeA-39137 |
| HUAS57 | Abasár-Bolt-tető | S57 | tooth | 686 | 14 | 1279-1303 (79%); 1368-1379 (16,4%) | INR Debrecen | DeA-39135 |
| HUAS59B | Abasár-Bolt-tető | S59B | rib fragment | 628 | 12 | 1300-1325 (43,7%); 1352-1395 (51,7%) | INR Debrecen | DeA-39138 |
| HUAS261 | Abasár-Bolt-tető | S261 | tooth | 692 | 13 | 1277-1301 (88,4%); 1371-1378 (7%) | INR Debrecen | DeA-39129 |
| HUAS262 | Abasár-Bolt-tető | S262 | tooth | 690 | 12 | 1278-1301 (87,8%); 1371-1378 (7,6%) | INR Debrecen | DeA-39130 |
| HUAS309 | Abasár-Bolt-tető | S309 | tooth | 608 | 13 | 1305-1365 (76,6%); 1383-1398 (18,9%) | INR Debrecen | DeA-39131 |
| HUAS341 | Abasár-Bolt-tető | S341 | tooth | 383 | 12 | 1455-1505 (78,8%); 1596-1618 (16,6%) | INR Debrecen | DeA-39132 |
| HUAS390 | Abasár-Bolt-tető | S390 | tooth | 858 | 13 | 1169-1222 (95,4%) | INR Debrecen | DeA-39133 |
| HUAS450 | Abasár-Bolt-tető | S450 | tooth | 618 | 12 | 1301-1329 (43,8%); 1341-1369 (29,3%); 1379-1396 (22,3%) | INR Debrecen | DeA-39134 |

*Notes:*

[1]Software and settings: OxCal 4.4.4, IntCal20

[2]Measuring labs: INR Debrecen: AMS laboratory of the Institute for Nuclear Research, Hungarian Academy of Sciences, Debrecen, Hungary

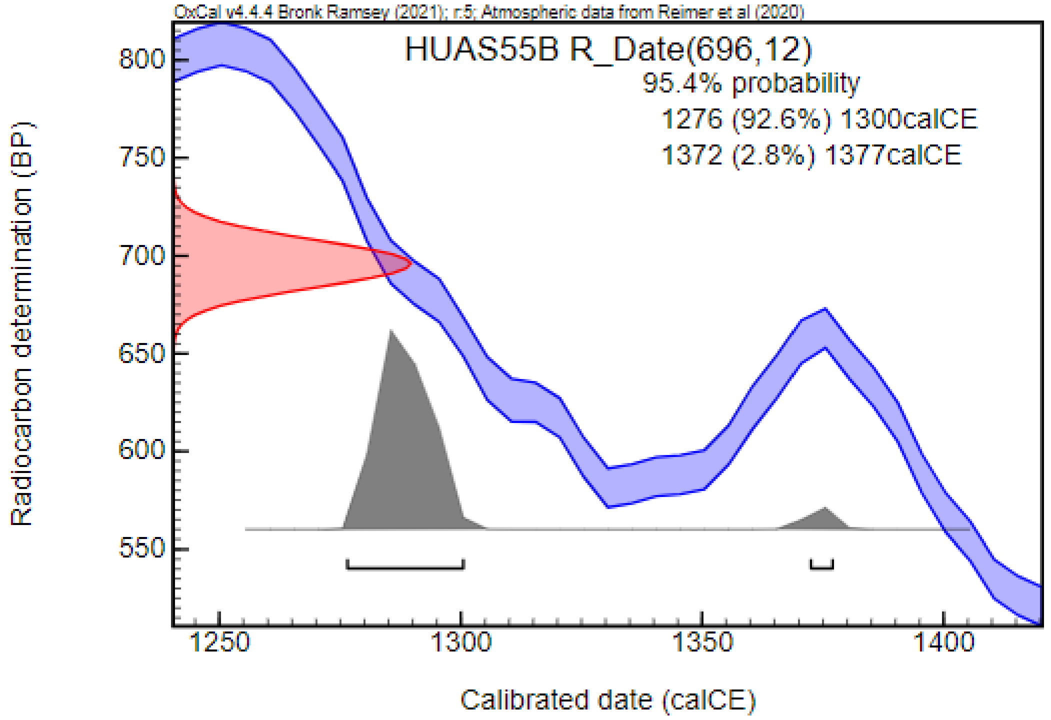
OxCal v4.4.4 Bronk Ramsey (2021); r:5; Atmospheric data from Reimer et al (2020)
HUAS55B R_Date(696,12)
95.4% probability
1276 (92.6%) 1300calCE
1372 (2.8%) 1377calCE
Radiocarbon determination (BP)
Calibrated date (calCE)

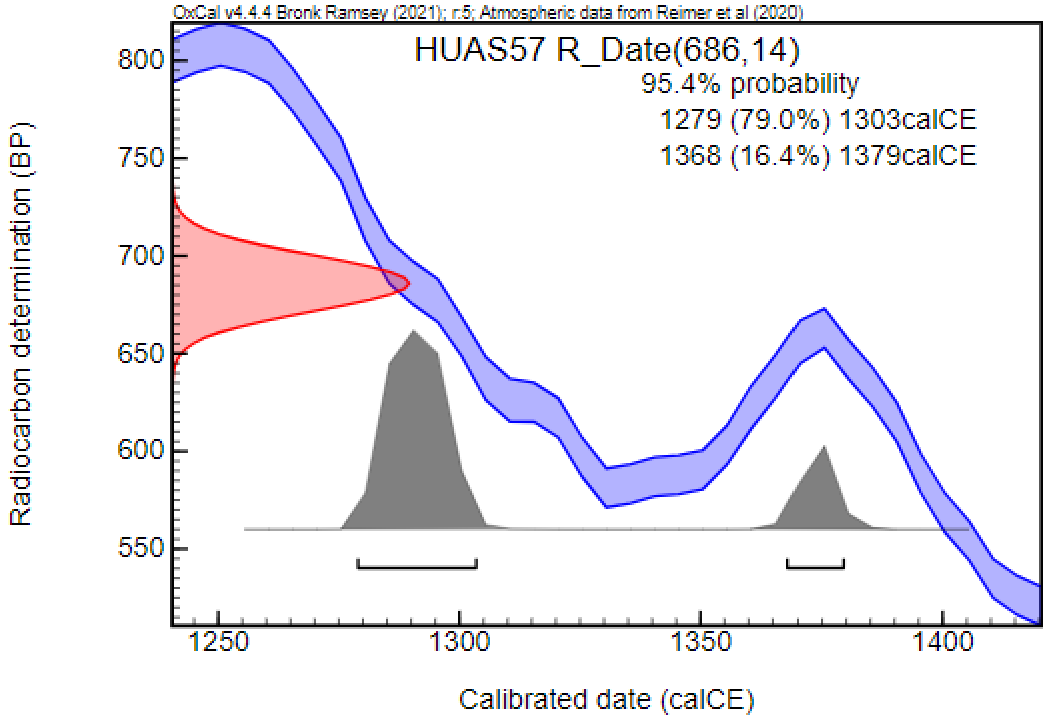
OxCal v4.4.4 Bronk Ramsey (2021); r:5; Atmospheric data from Reimer et al (2020)
HUAS57 R_Date(686,14)
95.4% probability
1279 (79.0%) 1303calCE
1368 (16.4%) 1379calCE
Radiocarbon determination (BP)
Calibrated date (calCE)

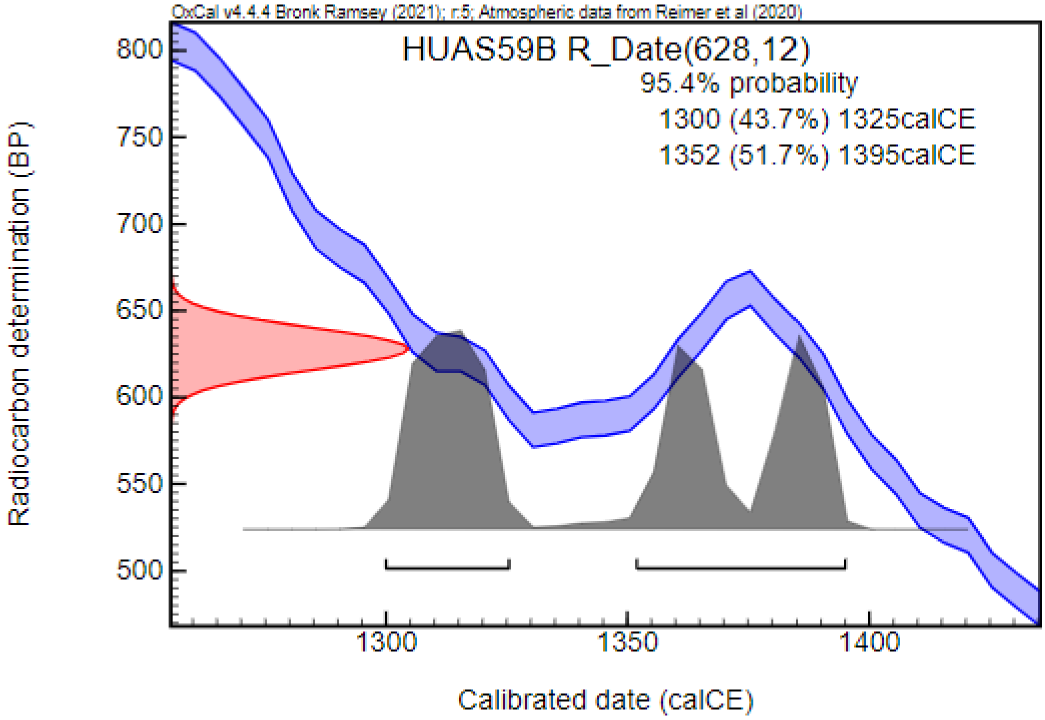
OxCal v4.4.4 Bronk Ramsey (2021); r:5; Atmospheric data from Reimer et al (2020)
HUAS59B R_Date(628,12)
95.4% probability
1300 (43.7%) 1325calCE
1352 (51.7%) 1395calCE
Radiocarbon determination (BP)
Calibrated date (calCE)

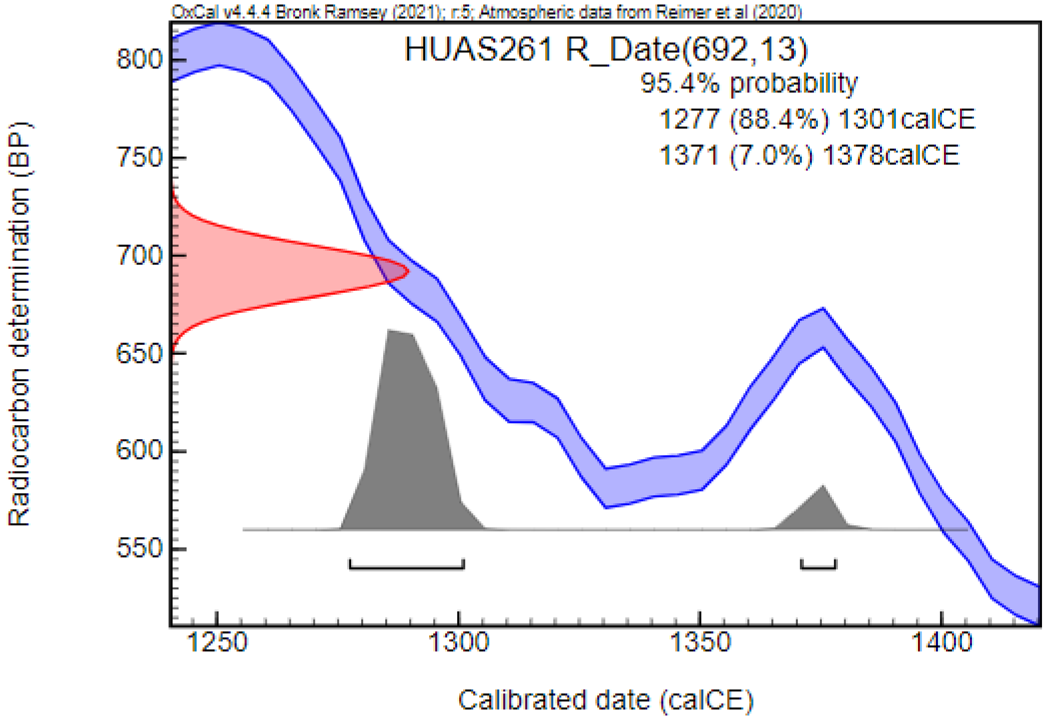
OxCal v4.4.4 Bronk Ramsey (2021); r:5; Atmospheric data from Reimer et al (2020)
HUAS261 R_Date(692,13)
95.4% probability
1277 (88.4%) 1301calCE
1371 (7.0%) 1378calCE
Radiocarbon determination (BP)
Calibrated date (calCE)

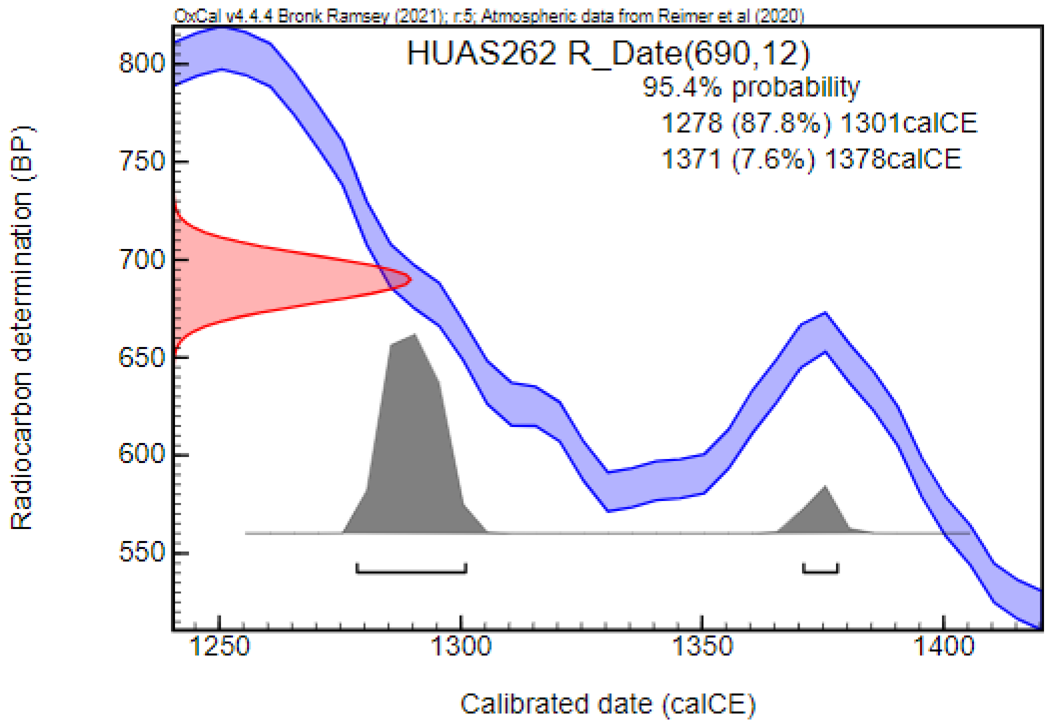
OxCal v4.4.4 Bronk Ramsey (2021); r:5; Atmospheric data from Reimer et al (2020)
HUAS262 R_Date(690,12)
95.4% probability
1278 (87.8%) 1301calCE
1371 (7.6%) 1378calCE
Radiocarbon determination (BP)
Calibrated date (calCE)

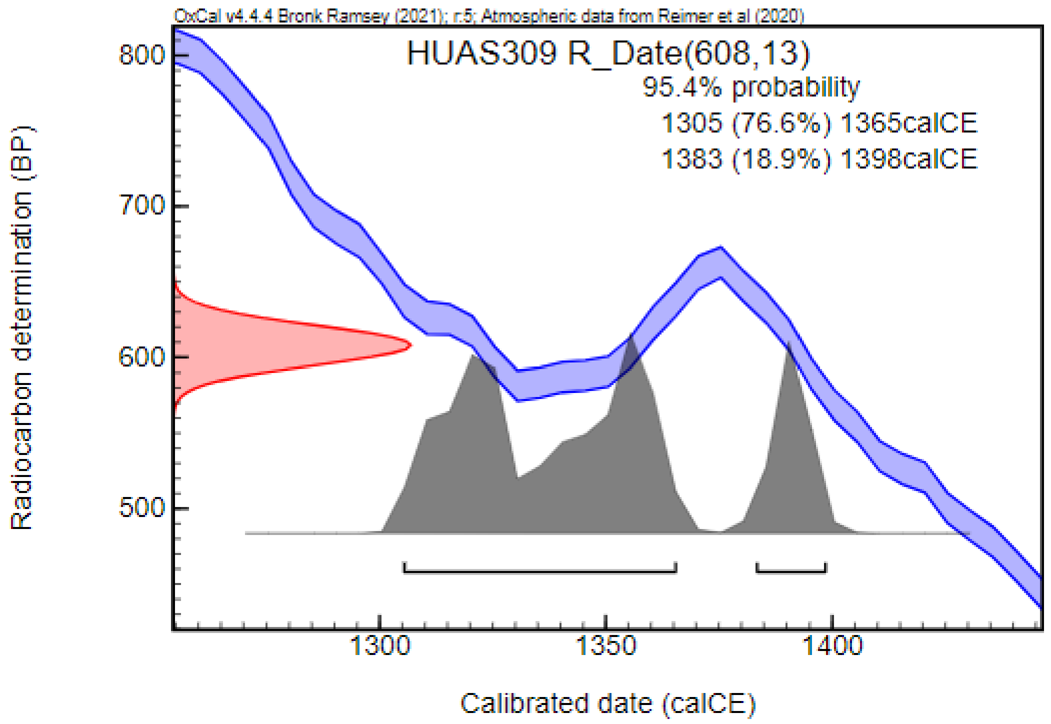
OxCal v4.4.4 Bronk Ramsey (2021); r:5; Atmospheric data from Reimer et al (2020)
HUAS309 R_Date(608,13)
95.4% probability
1305 (76.6%) 1365calCE
1383 (18.9%) 1398calCE
Radiocarbon determination (BP)
Calibrated date (calCE)

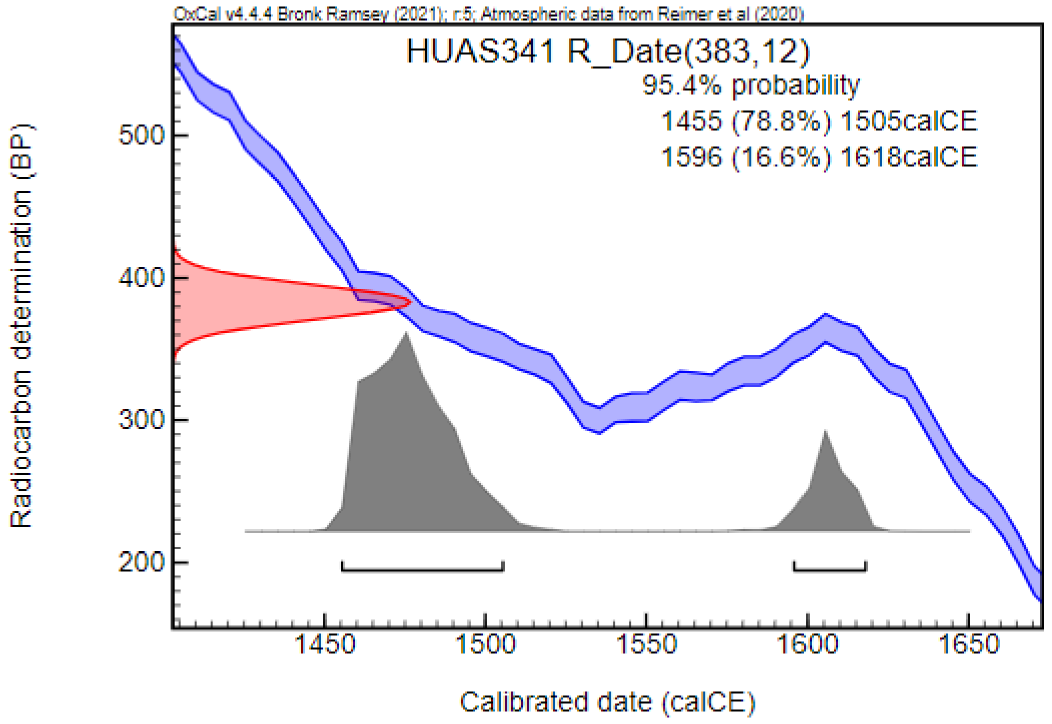

OxCal v4.4.4 Bronk Ramsey (2021); r:5; Atmospheric data from Reimer et al (2020)
HUAS341 R_Date(383,12)
95.4% probability
1455 (78.8%) 1505calCE
1596 (16.6%) 1618calCE
Radiocarbon determination (BP)
500
400
300
200
1450
1500
1550
1600
1650
Calibrated date (calCE)

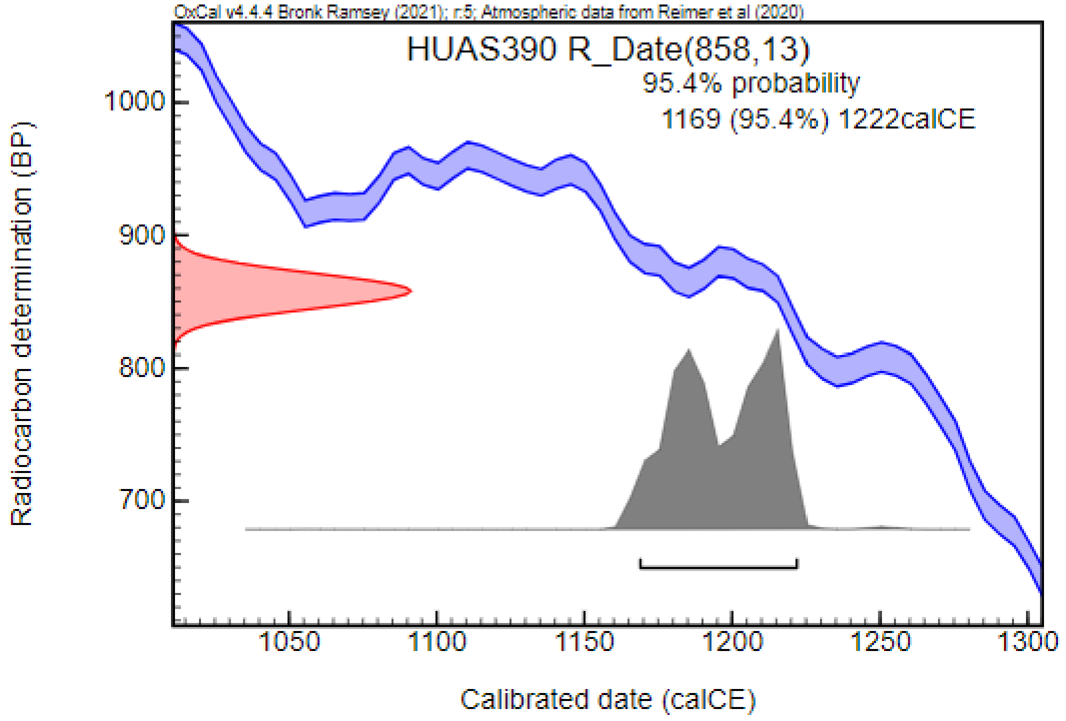

OxCal v4.4.4 Bronk Ramsey (2021); r:5; Atmospheric data from Reimer et al (2020)
HUAS390 R_Date(858,13)
95.4% probability
1169 (95.4%) 1222calCE
Radiocarbon determination (BP)
1000
900
800
700
1050
1100
1150
1200
1250
1300
Calibrated date (calCE)

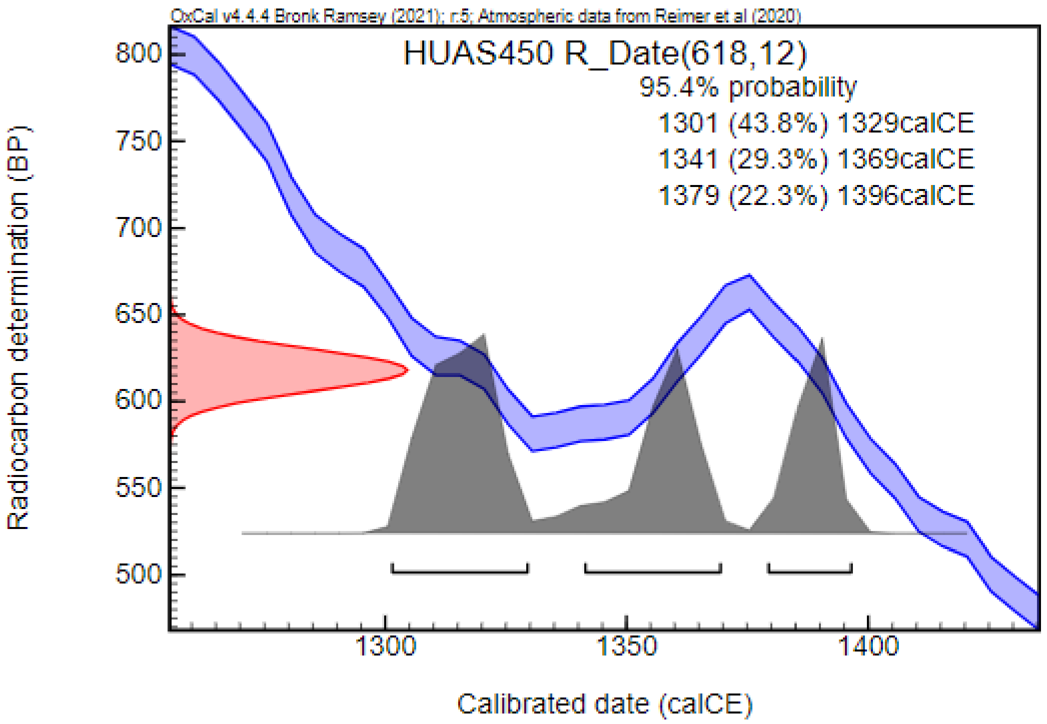
OxCal v4.4.4 Bronk Ramsey (2021); r:5; Atmospheric data from Reimer et al (2020)
HUAS450 R_Date(618,12)
95.4% probability
1301 (43.8%) 1329calCE
1341 (29.3%) 1369calCE
1379 (22.3%) 1396calCE
Radiocarbon determination (BP)
800
750
700
650
600
550
500
1300
1350
1400
Calibrated date (calCE)

**Extended figures**

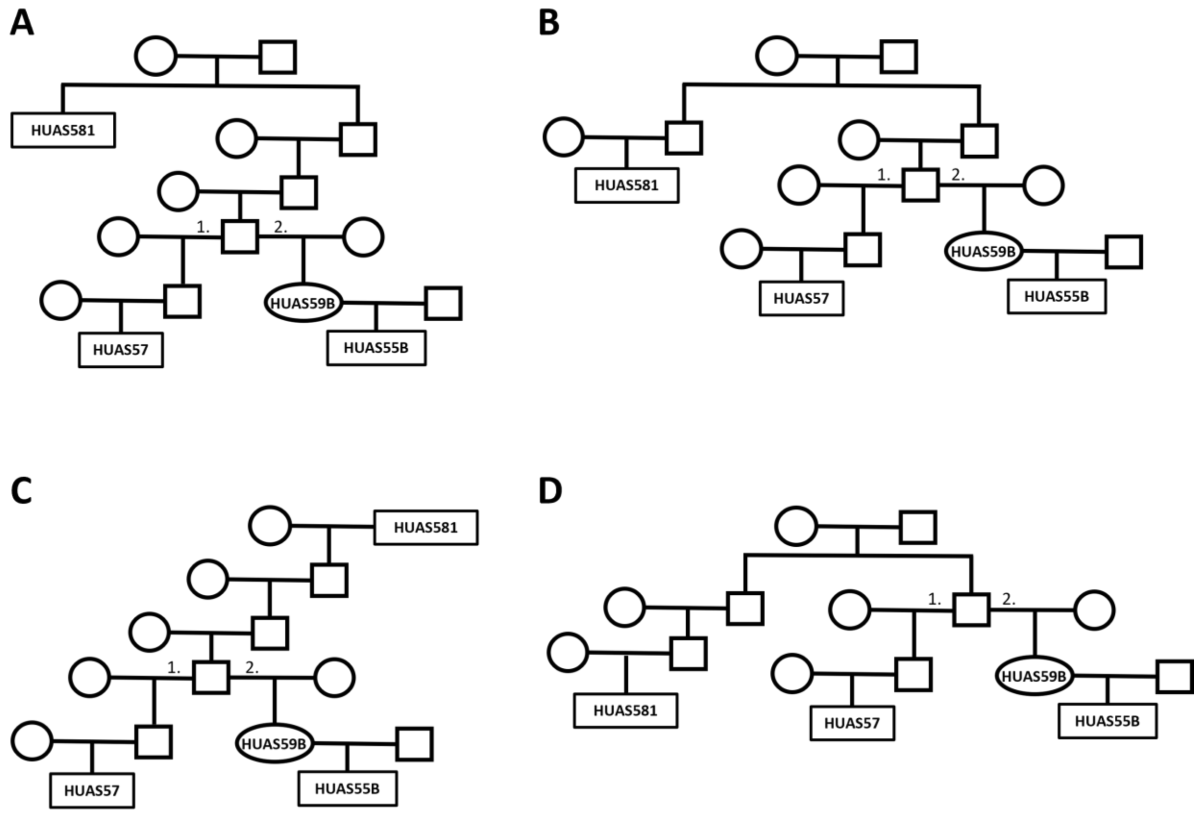

**Extended figure 1.** Reconstructed equivalent family trees of the larger family from Abasár Bolt-tető site based on the results with correctKin.

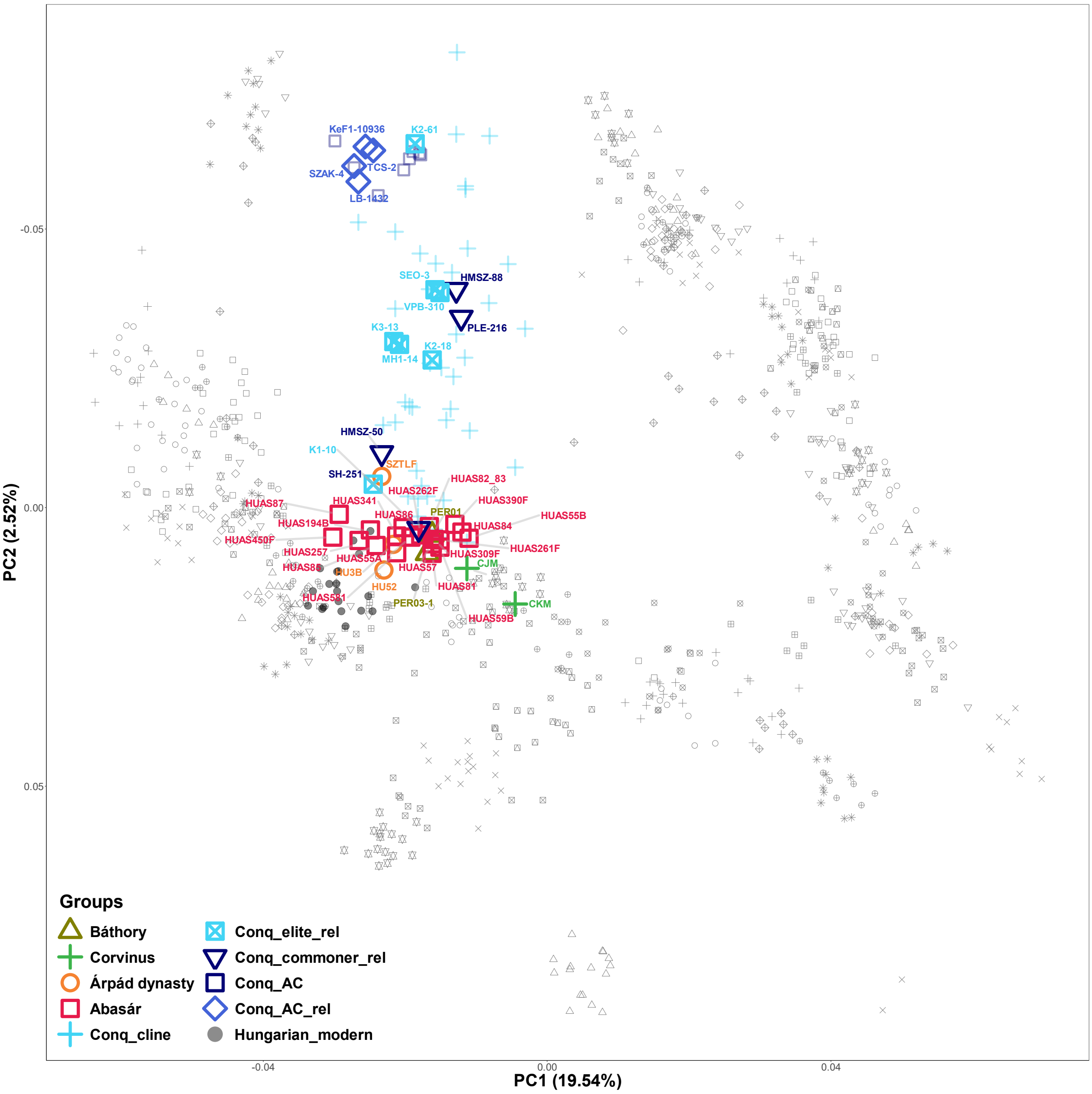

**Extended figure 2.** Figure 5 supplemented with sample IDs.

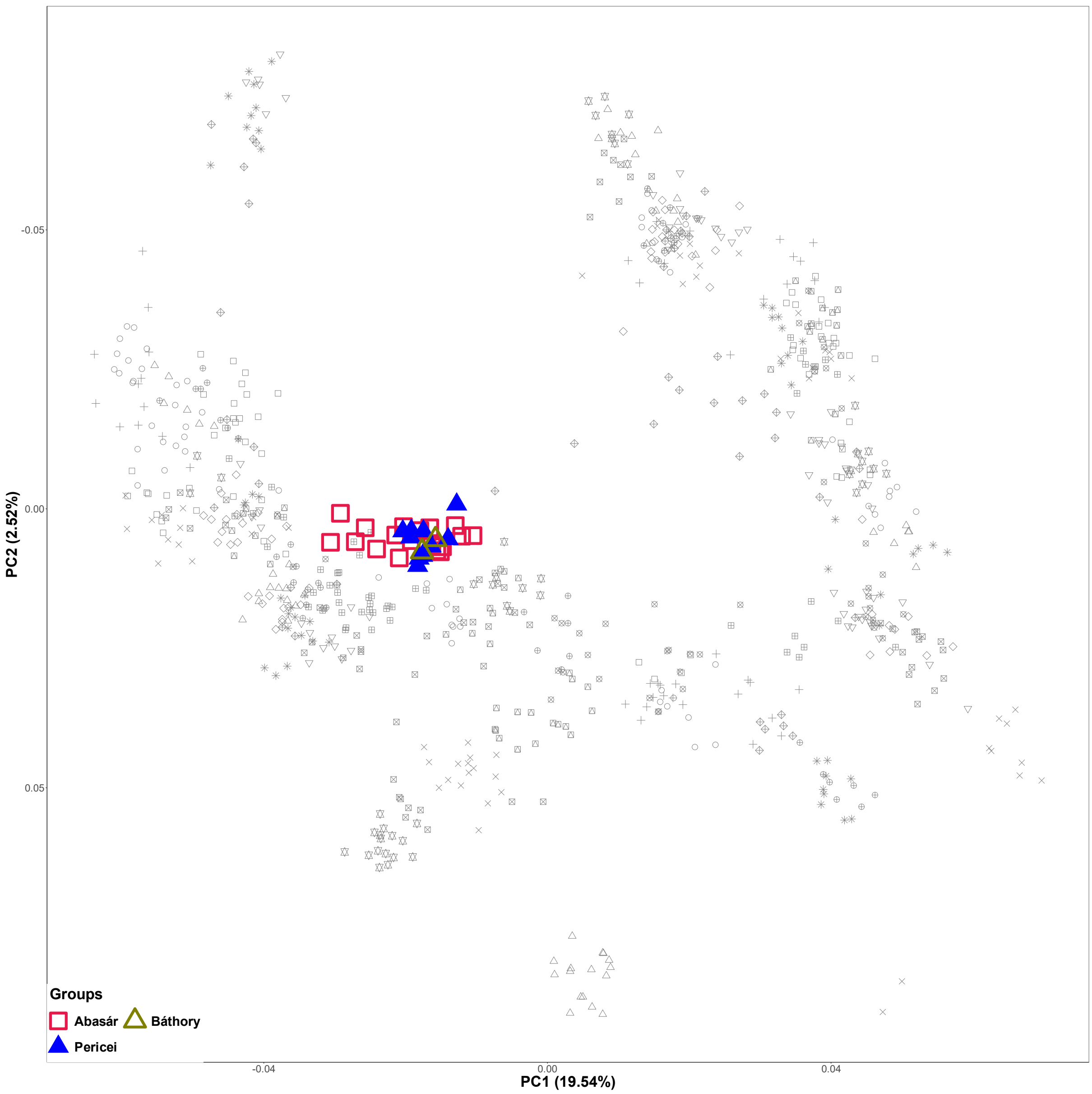

**Extended figure 3.** European PCA of Abasár samples (red squares) supplemented with the individuals from the Báthory cemetery at Pericei site (blue triangles) and the two Báthory family members (green triangles). The overlap between the two cemeteries indicates similar genome pattern for the two medieval Hungarian aristocratic group.

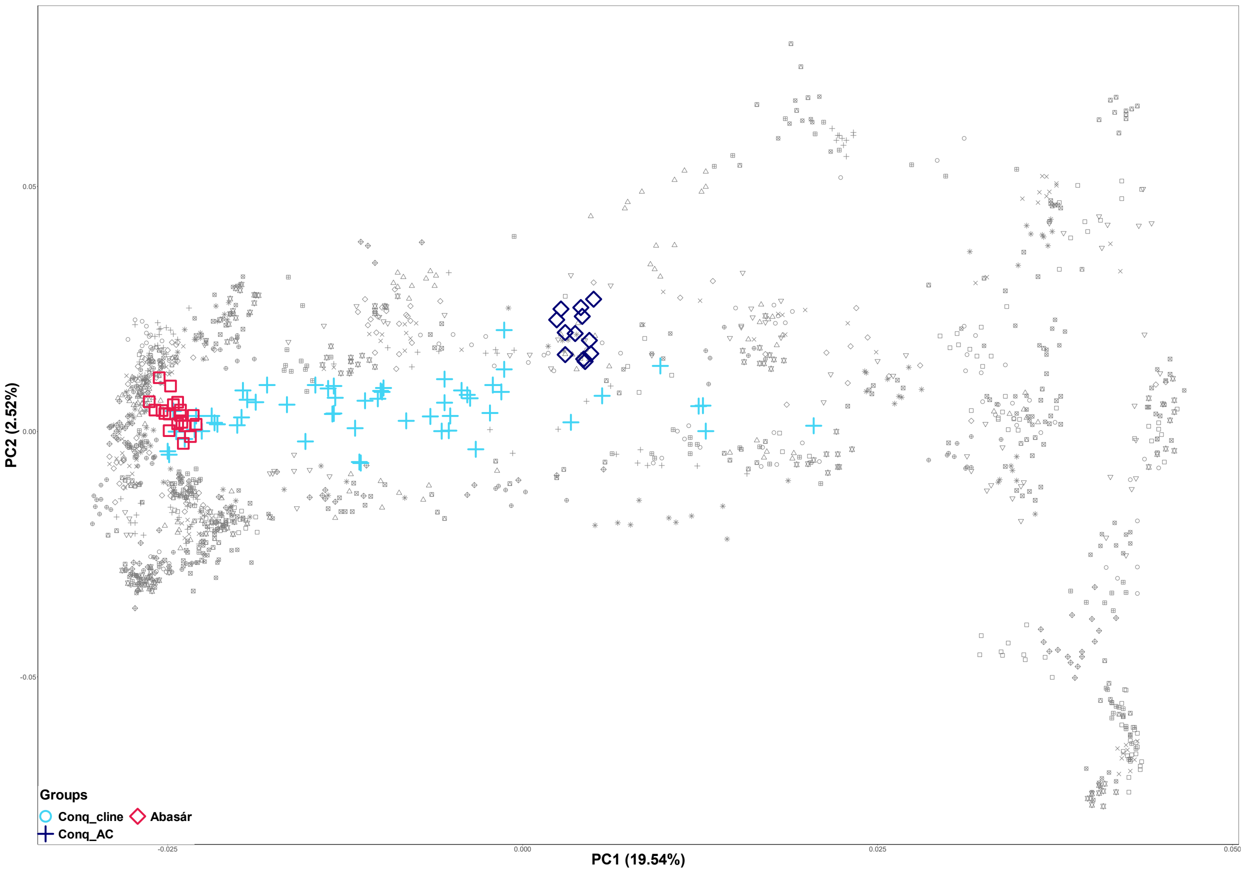

**Extended figure 4.** Eurasian_PCA of Abasár samples supplemented with the Conqueror cline (light blue crosses) and Conqueror Asia Core samples (blue squares). The cluster of Abasár samples overlaps with the Conqueror cline, indicating minor Asian patterns within their genomes feasibly originating from the conquerors.
